# Population dynamics of fruit flies (Diptera: Tephritidae) in a semirural area under subtropical monsoon climate of Bangladesh

**DOI:** 10.1101/2024.09.22.614336

**Authors:** Mahfuza Momen, Md. Aftab Hossain, Kajla Seheli, Md. Forhad Hossain, Md. Abdul Bari

## Abstract

Fruit flies belonging to Tephritidae family are highly destructive agricultural pests, posing a significant threat to various fruits and vegetables grown in Bangladesh. A comprehensive year-round survey was conducted at Atomic Energy Research Establishment (AERE) campus located in the central region of Bangladesh. Three types of male lures (methyl eugenol, cue-lure and zingerone) were used to detect and assess the diversity of pest fruit fly species. A total of seventeen species of Tephritidae fruit flies were detected in this survey. The *Bactrocera carambolae* fruit fly has been discovered for the first time in our survey area, indicating spread of its range towards the north-west region from its previous detection sites (Chattogram and Sylhet Divisions) in Bangladesh. Among the detected pest species, we identified six abundant species: *Bactrocera dorsalis, Zeugodacus cucurbitae, Zeugodacus tau, Bactrocera rubigina, Bactrocera zonata*, and *Dacus longicornis*. The most abundant species was the polyphagous fruit pest *B. dorsalis*, comprising 76.83% of all captured flies. The species *Z. cucurbitae* was the second most abundant, representing 13.82% of the total trapped flies. The fitted curve to survey data using Gaussian mixture model revealed the existence of overlapped subgroups in the population of *B. dorsalis* and *Z. cucurbitae*. In addition, our statistical analysis of the six abundant Tephritidae fruit fly species revealed correlation of population dynamics with several factors including temperature, rainfall, humidity, photoperiod, and fruiting time of host plant species in the selected area.

## 1. Introduction

Tephritidae fruit flies, comprising over 5000 species classified in 500 genera, are found widespread across the globe (Scolari *et al*., 2021). Most species in this group appear as destructive pests for horticulture crops, causing considerable damage to fruits and vegetables. As a result, the quality and quantity of marketable agricultural products reduces significantly. Ultimately, infestations of these fruit flies result in enormous annual economic losses in the agricultural industry. Due to the highly devastating impact on agriculture, these flies are recognized as significant quarantine pests. In response, many countries have implemented strict quarantine measures to protect their agriculture industries.

Because of high economic impact of fruit flies in the agricultural sector, researchers across the world have conducted significant biological and taxonomic studies on this group of fruit flies (Doorenweerd *et al*., 2018). Understanding the year-round population variation of a pest in relation to weather conditions is also crucial for developing a successful integrated pest management (IPM) program and regulatory rules for international trade. In addition, patterns of population fluctuations can provide valuable insights for farmers to develop effective pest management strategies well in advance. This proactive approach can lead to a reduction in crop losses, ultimately benefiting the agricultural industry.

Bangladesh, a part of Indian subcontinent, experiences a tropical monsoon climate marked by significant seasonal fluctuations in rainfall, warm temperatures and high humidity levels. Many abiotic (e.g. temperature, relative humidity, photoperiod, rainfall, wind, and other climatic conditions) and biotic (e.g. predators, parasitoids, host plant availability and host species richness) factors have a significant impact on the quantity and diversity of fruit flies throughout the year. Climate change also has a significant impact on the population dynamics and geographical distribution of fruit flies. Therefore, monitoring and collecting up-to-date information of fruit fly populations is crucial for any territory. However, very limited information is available about the population dynamics of fruit flies in this region.

To the best of our knowledge, the first taxonomic studies of fruit flies were reported by Fabricius in 1794 in the Indian subcontinent (including Bangladesh region) where three fruit flies were detected (Agarwal & Sueyoshi, 2005). Despite their economic importance, the detection of fruit flies progressed slowly in this region during the nineteenth century. More comprehensive and systematic scientific surveys began much later. In the first half of the twentieth century, Senior-White reported 87 fruit fly species of undivided India (Kapoor, 1970). The same researcher also reported fruit flies from east Pakistan (current name Bangladesh) in 1922 (Kapoor et al., 1980; Senior-White, 1922, pp. 83-169). A total of 11 fruit fly species were detected in the Bangladesh region during 1767 to 1951 (Kapoor et al., 1980). According to a recent survey by Leblanc *et al*., presence of 37 Tephritidae fruit fly species was reported in Bangladesh (Leblanc *et al*., 2019).

However, previous studies on fruit flies provided very limited information on the correlation between environmental factors (e.g. photoperiod) and population dynamics in this territory.

In a region with tropical climate like Bangladesh, a wide range of host plants provides ideal breeding conditions for fruit flies with prolonged breeding season, allowing them to reproduce multiple times a year. Therefore, many major pest fruit flies in this region are multivoltine (Choudhary *et al*., 2019). Information on the presence of coexisting overlapping generations or subgroups is also crucial for understanding population dynamics for pest management. However, previous studies on fruit fly in this region did not focus on the existence of the subgroups of fruit fly population.

This study presents a year-round comprehensive survey on Tephritidae fruit fly in the area of Atomic Energy Research Establishment (AERE) Campus, Dhaka, Bangladesh. Along with the evaluation of species detection range of the three types of male lures used in the survey, our study presents a detailed statistical analysis of the diversity of Tephritidae fruit fly species using species richness index (Shannon diversity index and iChao1 estimator) and species accumulation curve. In our study, we also analyzed the population dynamics of the six most abundant fruit fly species in relation to various environmental factors including temperature, humidity, photoperiod, rainfall, and the fruiting stage of host plants. By examining these correlations, we were able to uncover significant correlations that revealed the relationships between the fruit fly populations and their surrounding ecosystem. Moreover, for the first time, we identified the existence of subgroups of the two major abundant fruit fly species *Bactrocera dorsalis* (Hendel) and *Zeugodacus cucurbitae* (Coquillett) population in this area using Gaussian mixture model. By identifying dominant Tephritidae fruit fly species and quantifying their abundance levels and population dynamics, this research provides crucial insights for developing effective pest management strategies for Bangladesh.

## 2. Materials and Methods

### 2.1. Survey area

Our selected survey area was Atomic Energy Research Establishment (AERE) situated in the central region of Bangladesh, at coordinates 23° 57’ 18.72”N and 90° 17’ 30.39”E (**Fig. 1**). The monitoring survey spanned a full year from October 2020 to September 2021 and focused on the uncontrolled Tephritidae pest populations present in the AERE campus. This semirural area contains a diverse range of plants (over 400 plant species), such as trees, wild shrubs, and crop plants. Additionally, there were numerous small water bodies scattered throughout the area. Moreover, human dwellings were also included in this area, creating a more complex habitat that supports a wider variety of fauna in this area. In AERE area, there was a representation of a wide variety of major crops, fruits, and vegetables cultivated in Bangladesh. Frequent fruit fly infestation was also recorded in this area. Therefore, this area was an ideal representative for most semi agroforest areas to study the diversified Tephritidae pest populations for the application of sterile insect technique.

**Fig. 1:**
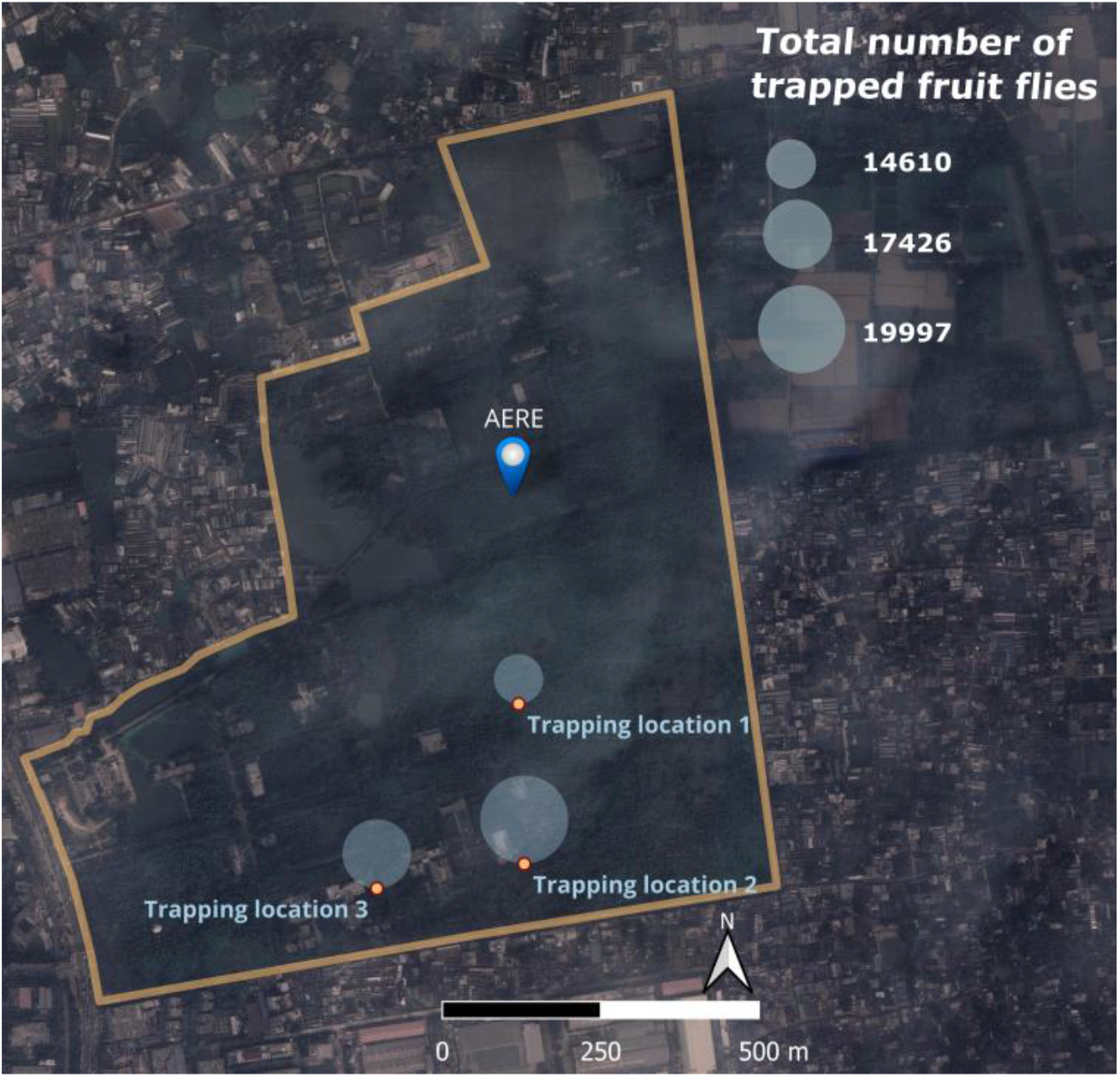
Landmark position of the three trapping sites at Atomic Energy Research Establishment (AERE) area: trapping location 1 (site 1), trapping location 2 (site 2) and trapping location 3 (site 3). At each trapping site, three types of lure traps (ME, CL, and Zn) were placed ten meters away from one another. The relative radius of the trapping locations represents the comparison of the total number of fruit fly trapped in that location. The figure was generated with QGIS (https://qgis.org/), version 3.40. The map of the study area was sourced from Google Earth.

### 2.2 Trapping method for fruit fly detection

Three different types of male lure-based traps were used to conduct an annual survey of a wide variety of Tephritidae fruit flies in the study area.

The trapping materials used for this study include:

- Attractants: Methyl eugenol (ME), cue-lure (CL) and zingerone (Zn)
- Insecticide: Insecticide strip containing 10% dichlorvos (2,2-dichlorovinyl dimethyl phosphate)
- Preservative: Liquid 10% propylene glycol (propane-1,2-diol)
- Devices for trapping: Modified Lynfield trap consists of a plastic container (Globe scientific specimen collection container, 120 ml) with a blue lid, an overhanging shade and two entrance holes.

Globe scientific specimen collection containers (6.35 cm diameter and 7.62 cm height) were used to make these traps. Two circular (1 cm diameter each) lateral holes were drilled below the screw cap to create fly entry points. Commercially available male-specific lure methyl eugenol (ME) and cue-lure (CL) 2 cm plug (Scentry Biologicals, Billings, Montana, USA) containing 2 g of lure were used as attractants for trapping Tephritidae fruit flies. In addition, comparatively new male lure Zingerone (Zn) (Sigma-Aldrich, St. Louis, Missouri, USA) dipped 2 cm cotton dental wick was also used as an attractant for trapping Tephritidae fruit flies. Controlled-release formulations were utilized for these volatile lures to prolong the effectiveness of attractants in the field.

#### 2.2.1. Killing and preserving agents

An insecticide strip (10 ×15 mm) containing 10% dichlorvos (Vaportape II™, Hercon Environmental, Emingsville, Pennsylvania, USA) was placed inside the trap to knock down and kill attracted male fruit flies that entered the trap. In order to slow the drying of the attractant and preserve captured flies, 60 ml of 10% propylene glycol solution was added to the trap.

The wire holding the trap was coated with TangleTrap glue (Tanglefoot Company, Grand Rapids, Michigan, USA) to keep the captured flies safe from ants and other predatory threats. A disposable plastic plate with a diameter of 15 cm was placed over the trap to prevent the rainwater from getting inside.

#### 2.2.2. Trap deployment

Traps were placed within the southern agroforest area of AERE, where no pest control measures had been implemented. Traps were baited with one of the male lures (methyl eugenol, cue-lure or zingerone) and placed in trees 1.8 meters above the ground. Traps were placed in partially shaded areas of plants located near fruit fly nesting and feeding spots.

All the traps were spatially distributed in three suitable locations using global positioning system (GPS) to maximize the chances of detecting the expected pest fruit fly population (**Fig. 1**). In each trapping location, three different lure (ME, CL and Zn) traps were placed at a minimum distance of 10 meters away from each other. The positions of the traps were georeferenced using a GPS tracker device.

#### 2.2.3. Trap servicing and inspection

We serviced (i.e. rebaiting) all traps every six weeks to ensure they stayed clean and operated properly. During servicing, extra care was taken to prevent mixing different attractant types (CL, Zn, and ME) and avoid cross-contamination between traps. Separate gloves were worn when changing each lure, and previous lures were disposed of away from the traps. Inspection intervals for sample collection were adjusted to seven days according to the prevailing environmental conditions and pest situations. During the inspection, specimens were collected in vials containing 95% ethanol and labeled with trap information details (GPS coordinates, attractant type and date).

Captured flies were then identified and counted by skilled taxonomists at the research laboratory of Insect Biotechnology Division, Institute of Food and Radiation Biology, AERE. Identification was made based on the morphological characteristics of the collected specimen using a stereo microscope (Leica DMC2900) and taxonomic keys described by Drew and Romig (Drew & Romig, 2013).

#### 2.2.4. Trapping record

Microsoft Excel spreadsheet was used to maintain accurate trapping records for precise data tracking and analysis. It included detailed information such as location of trap (GPS coordinates), type of attractant used, dates of servicing and inspection, as well as the taxonomic identity of captured fruit flies along with their corresponding quantity.

Data of the weekly captured six abundant pest fruit fly species (*Bactrocera dorsalis, Zeugodacus cucurbitae, Z. tau, B. rubigina, B. zonata, and Dacus longicornis)* (**Fig. 4**) were analyzed to determine their temporal distribution throughout one year trapping period.

FTD (Flies per trap per day) was used for various types of analysis, including temporal changes in fruit fly populations. FTD is a population index that indicates the average number of flies of the target species captured per trap per day during a specified period in which the trap was exposed in the field (IAEA, 2003).

FTD was determined by calculating the number of fruit flies captured in a trap each day, using the following formula:

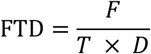

Where, *F* is the total number of fruit flies, *T* is the number of inspected traps, *D* is the average number of days of exposed traps in the field between two inspections.

### 2.3 Data analysis and statistical methods

#### 2.3.1. Shannon diversity index

We used the Shannon diversity index (*H*) (**Fig. 5**) to measure the diversity of the Tephritidae fruit fly species richness and abundance in three sites of AERE campus. The bias-corrected maximum likelihood estimation (MLE bc) estimator for Shannon diversity index given by Chao and Shen (Chao & Shen, 2003) was used to calculate this index. The following formula was used for this calculation:

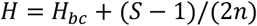

where,

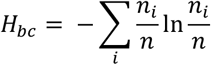

Here, *S* is the number of taxa, *n* is the total number of fruit flies, *ni* is the number of fruit flies of taxon *i*, and *H*_*bc*_ represents computed with a bias correction value.

#### 2.3.2. iChao 1

An estimate of species richness (*S*_*ichao*1_) was calculated based on iChao1 estimator (**Fig. 5**), which is an improved Chao1 estimator (Chiu *et al*., 2014).

We used bias correction version of Chao1 (an estimate of total species richness) (Chao, 1984) calculated by the following formula:

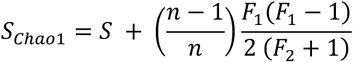

where, *F*_1_ is the number of singleton species and *F*_2_ the number of doubleton species. From Chao1 version with bias correction, we calculated *S*_*ichao*1_ while taking into account the numbers *F*_3_ and *F*_4_ of species observed 3 and 4 times (Chiu *et al*., 2014). The following formula was used for this calculation:

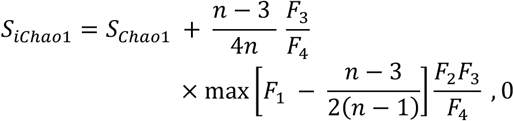

If *F*_4_ = 0, we used *F*_4_ = 1 to avoid division by zero.

#### 2.3.3. Species accumulation curve

In order to evaluate and compare diversity levels among the three trapping sites, we used species accumulation curves to estimate the expected number of observed species. We analyzed this data using Mao’s tau and standard deviation. To represent the data graphically (**Fig. 5**), we converted standard errors into 95% confidence intervals (Colwell *et al*., 2004).

#### 2.3.4. Gaussian mixture model

A Gaussian distribution (also called normal distribution) is a continuous probability distribution that is characterized by its symmetrical bell-curve (Jayakumari et al., 2023; Motulsky, 2018). Gaussian distribution has two parameters, and the probability density function can be represented by the following equation:

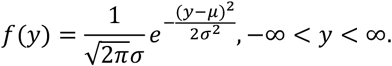

Where, *μ* is the mean and *σ* is the standard deviation. The bell curve is centered around *μ*.

Trap-captured insect population distribution as a function of time can be approximated to a bell-curve of Gaussian distribution or mixture of Gaussian distributions (Barclay et al., 2016; Lan & Chandran, 2011; Onufrieva & Onufriev, 2018; Soulsby & Thomas, 2012). Gaussian Mixture Model (GMM) is a probabilistic model that describes the distribution of a data set as the mixture of a finite number of local Gaussian distributions called Gaussian components, each representing a distinct cluster (bell-curve). In a Gaussian Mixture Model (GMM), linear combination of distinct or overlapped bell-curves can represent more complex population distribution (Reynolds, 2009; Van Os, 2022). By analyzing the composition of the overlapping clusters (bell-curve), entomologists can gain valuable clues regarding the coexistence of generations, the number of instars of a species (Barclay et al., 2016; Wu et al., 2013) etc. in a complex environment. If the survey duration for fruit flies spans more than one generation, during which oviposition and mortality occur over extended periods, the mixture model is better suited (Barclay et al., 2016; Manly, 1974; Wu et al., 2013). Therefore, population data fitting was performed using Gaussian mixture model. This enables the detection of population subgroups, which helps to make more precise forecasts and well-informed pest management decisions. All the statistical analyses were carried out by OriginPro^®^ software.

## 3. Results and Discussion

### 3.1. Species detection range of different parapheromone traps

The life cycle of Tephritidae fruit flies has two active phases (larva and adult) that inhabit ecologically distinct habitats, which makes it challenging to study how their populations change over time. Although biotic and abiotic variables influence both stages differently, pheromone traps are specialized devices for capturing their active adult stage to monitor the population trend of fruit flies in an area. These types of traps with a lure are an easy way to capture large numbers of pest flies in a short period of time. Current fruit fly control, monitoring, and detection programs utilize methyl eugenol (ME) and cue-lure (CL) (Doorenweerd *et al*., 2018; Leblanc *et al*., 2019; Vargas *et al*., 2010).

ME and CL are both strong lures designed specifically for attracting male flies (IAEA, 2003). In addition, Zingerone (Zn), a relatively recent addition to the category of male lures, has demonstrated an ability to attract several species of fruit flies, including those unresponsive to CL or ME (Doorenweerd et al., 2018; Tan & Nishida, 2000). It was discovered that this lure is highly attractive to numerous pest species of Bangladesh (Leblanc *et al*., 2019). These types of monitoring trapping can also help to locate clusters of Tephritidae and identify suitable locations for a target-specific control approach. Unfortunately, no other viable and sustainable option exists for monitoring both male and female Tephritidae fruit flies.

To assess the effectiveness and compare the detection range (**Fig. 2**) of parapheromone traps baited with CL, ME, or Zn, we maintained the traps for one year (October 1, 2020 to September 30, 2021) at three locations (**Fig. 1**) within our study area. The study revealed that the zingerone (Zn) lure was significantly more effective in attracting a wider range of fruit fly species at AERE compared to other lures (CL and ME). Specifically, traps baited with Zn detected eleven different species, whereas traps baited with CL and ME detected only nine and four species, respectively (**Fig. 2**). These findings emphasize the potential of Zn lure as an effective tool for detecting a wider range of pest species in an area.

**Fig. 2:**
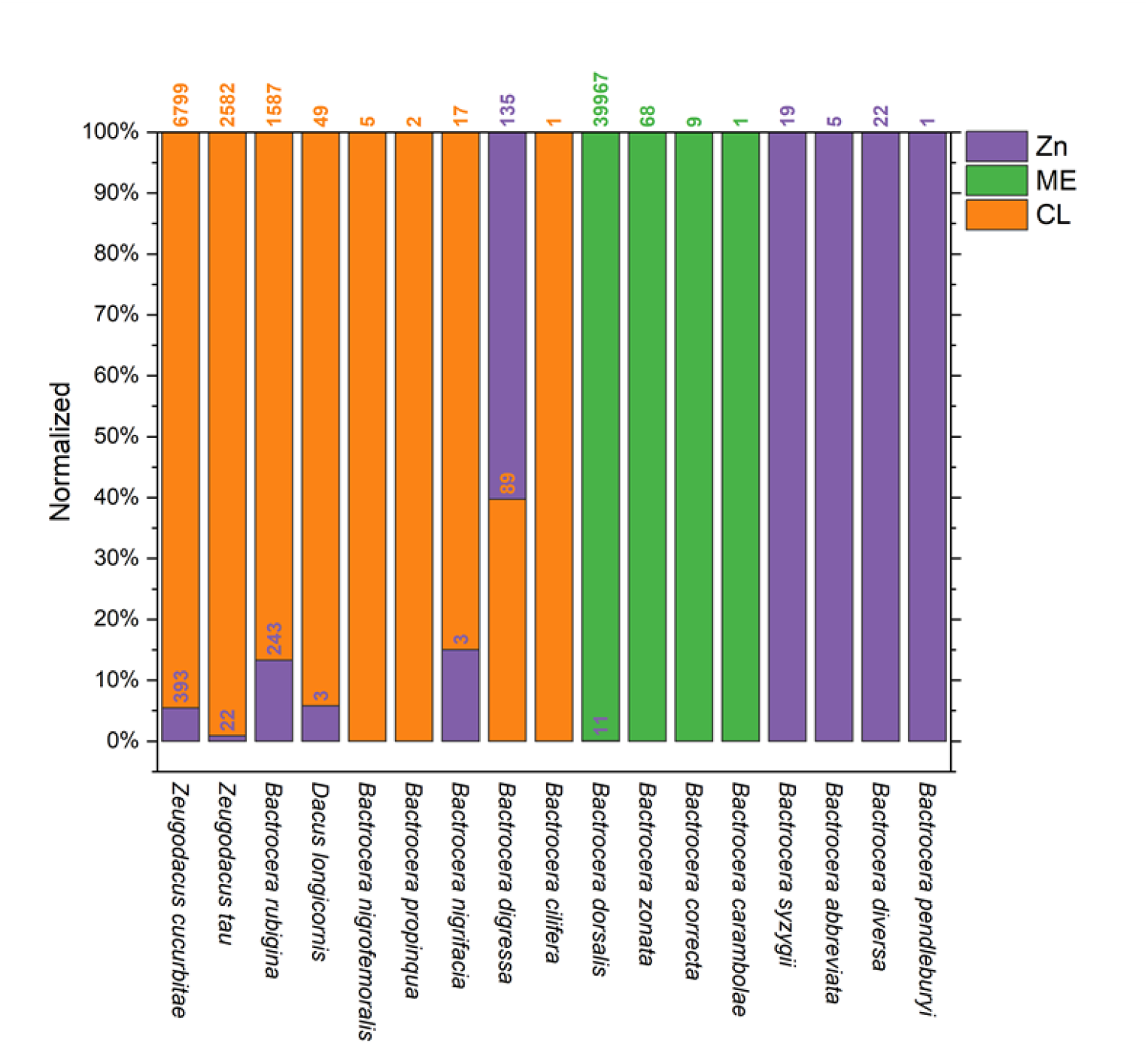
Detection Range of three types of male lures: cue-lure (CL), methyl eugenol (ME) and zingerone (Zn) for detected 17 Species of Tephritidae fruit flies trapped in AERE campus.

However, the small sample size of the detected fruit fly species by Zingerone (Zn) lure was inadequate to accurately detecting the population trend in our survey area (**Fig. 2**). Therefore, fruit flies that were caught in traps containing CL or ME lures were recorded to track the trend in population fluctuation of the six major fruit fly species. During our trapping period, we captured 52,033 Tephritidae fruit flies.

Among these Tephritidae fruit flies, we identified seventeen Tephritidae fruit fly species (**Fig. 2**). Out of seventeen species, six species were found to be abundant. The most abundant species was polyphagous fruit pest *B. dorsalis* (76.83% of all trapped flies); the second most abundant species was *Z. cucurbitae* (13.82%). *Z. tau* (5%) and the non-pest *B. rubigina* (3.52%) were in third and fourth positions, respectively (**Fig. 3**).

**Fig. 3:**
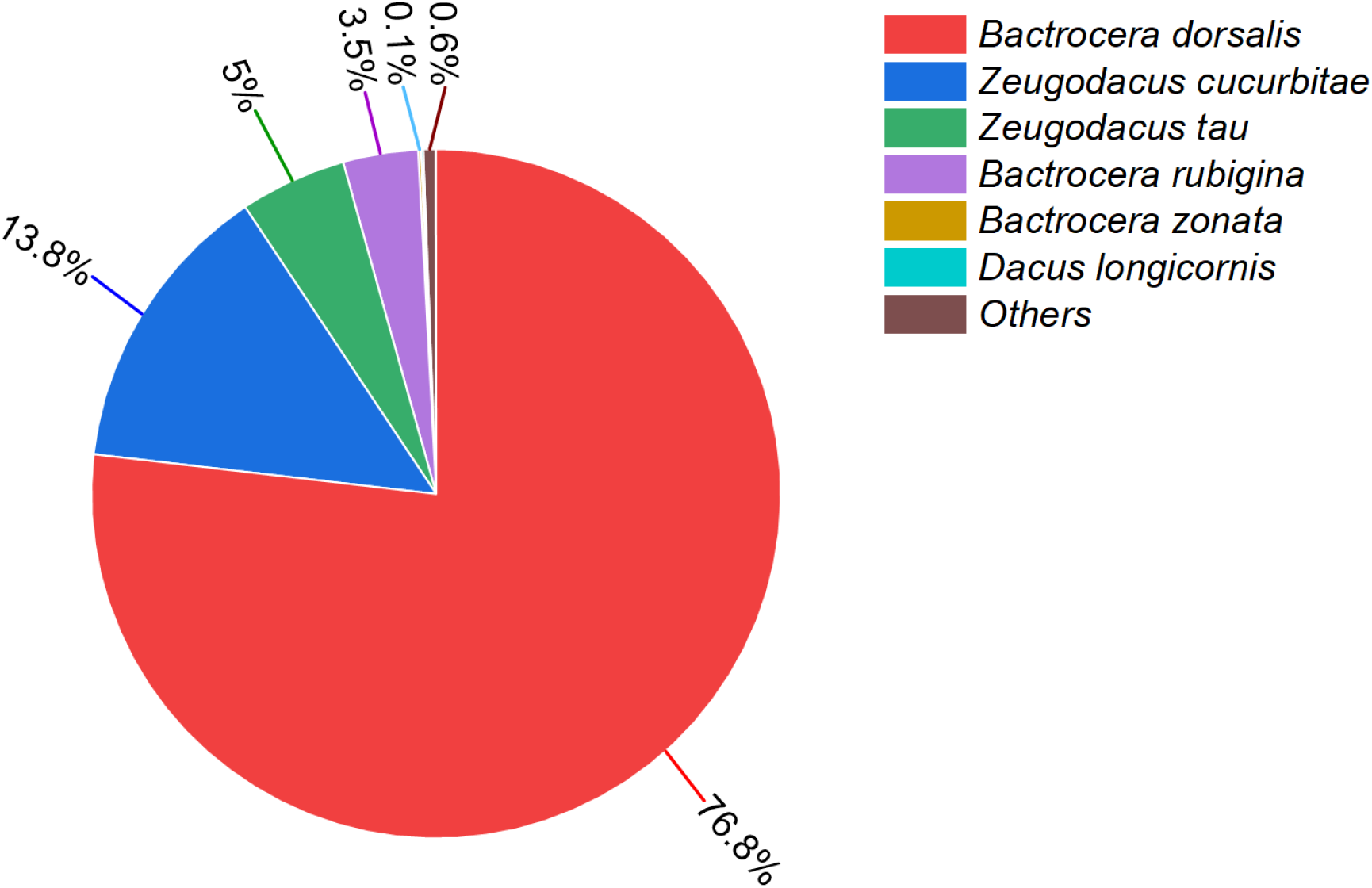
The percentage of the top six abundant Tephritidae species out of the seventeen detected species.

The other two major pest species, gathered in considerably lesser quantities (less than 0.25%), were polyphagous fruit pest *B. zonata* (0.13%) and cucurbit pest *D. longicornis* (0.10%). A similar dominance hierarchy pattern of detected fruit flies was also documented by Hossain *et al*. in a previous survey conducted in this area (Hossain *et al*., 2019). Surprisingly, we also detected a highly invasive fruit fly species *B. carambolae* in our survey area for the first time. This species was not detected before in this central region of Bangladesh where our survey area located. However, there was a report by Leblanc *et al*. of the initial detection of *B. carambolae* from surveys conducted from 2013 to 2018 in the Chattogram and Sylhet Divisions of Bangladesh (Leblanc *et al*., 2019). The discovery of *B. carambolae* during this survey indicates a notable spread of this species towards the north-west regions from its previously known detection sites of Bangladesh. This species has the potential to spread into other regions of the country. Therefore, it is crucial to regularly monitor the population of this pest in order to prevent rapid dissemination in invaded regions.

### 3.2. Species richness and diversity

Trapping in the three locations revealed varying species diversity and population sizes. The site 1 had sixteen species with 14,610 individuals, site 2 had twelve species with 19,997 individuals, and site 3 had fourteen species with 17,426 individuals (**Fig. 1**). This data highlights the diversity and distribution of species across the surveyed sites. It is evident that the dominance index and diversity index are inversely proportional. Therefore, the dominance index value is high at site 1, with only a few species being predominant in comparison to other sites (**Fig. 5**).

As we know, individual rarefaction estimates how many taxa are expected in a sample. Upon analyzing the individual rarefaction curve, it becomes evident that site 1 has the highest potential for discovering the maximum number of taxa followed by site 3. In contrast, the rarefaction curve for site 2 reaches a plateau, indicating that the sampling effort was sufficient, and the number of species detected using these lures was approaching near the actual total number of species existing at that site (**Fig. 5**). These local patterns of richness observed in the fruit fly community could be influenced by several factors, including their free flying abilities and other factors affecting their distribution. Conducting further research on the behavior and ecology of fruit flies could provide valuable insights into these localized richness patterns.

**Fig. 4:**
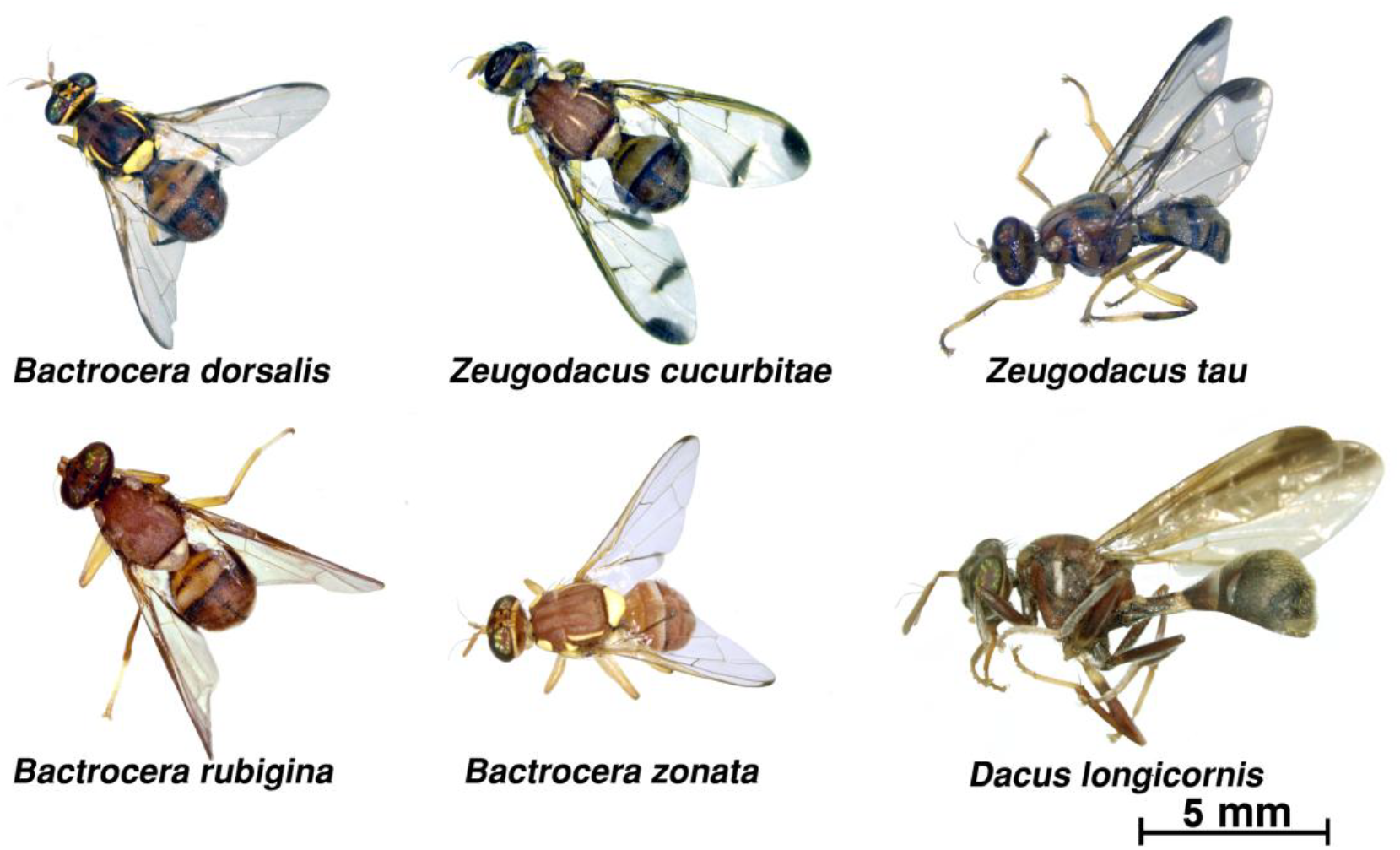
Pictures of the detected six topmost abundant tephritid fruit flies (*Bactrocera dorsalis, Zeugodacus cucurbitae, Zeugodacus tau, Bactrocera rubigina, Bactrocera zonata* and *Dacus longicornis*) in the AERE area.

**Fig. 5:**
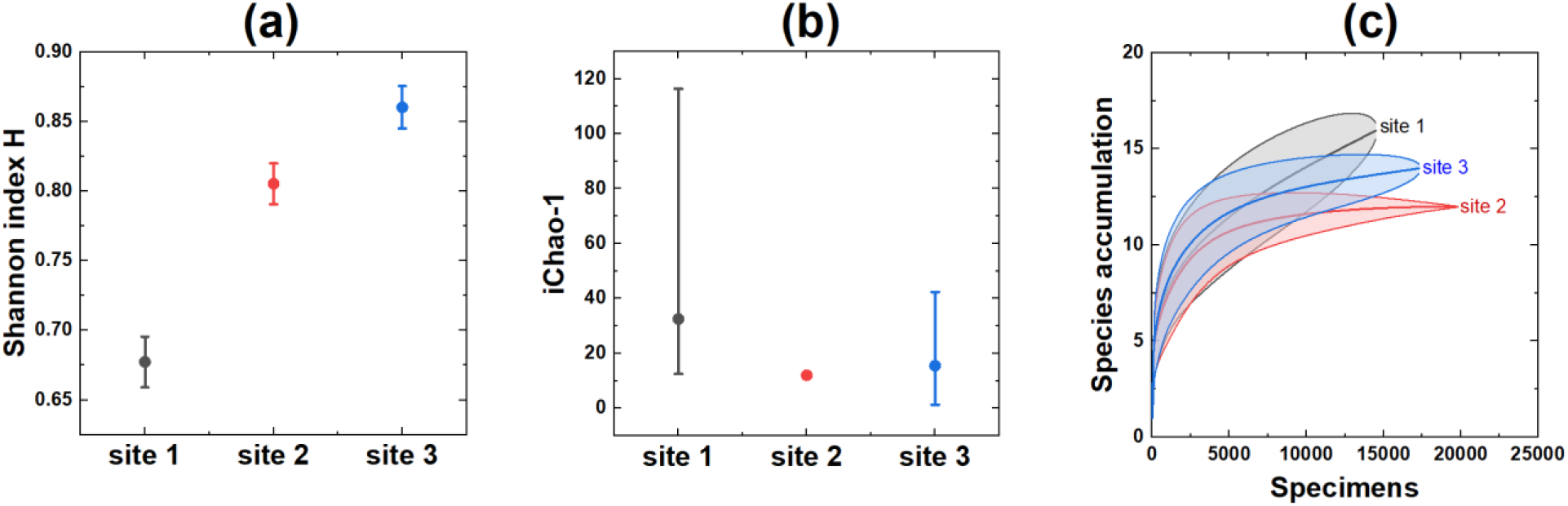
Species diversity index **(a)** Shannon index H, **(b)** ichao-1 and **(c)** rarefaction curve for the Tephritidae fruit fly community of AERE (Standard error is converted to 95% confidence level) of three trapping locations site 1, site 2 and site 3.

### 3.3 Month-wise detected Tephritidae fruit fly species

Tephritidae species and the months during which they were detected are illustrated in Figure 6 (**Fig. 6**). According to this study, large number of fruit fly species (ranging 11 to 14 Spp.) were most abundant from May to August. The highest number of species (14 Spp.) was detected in July. The lowest number of species (5 Tephritidae fruit fly species) was recorded in October. The number of species remained at or above 8 for the rest of the year. Population dynamics of these Tephritidae fruit fly species were influenced by seasonal variations of weather conditions and host availability (**Table 1**). In the next section, the relationship of population dynamics with weather conditions and host plant fruiting time are presented for the most abundant six Tephritidae species.

**Table 1.**
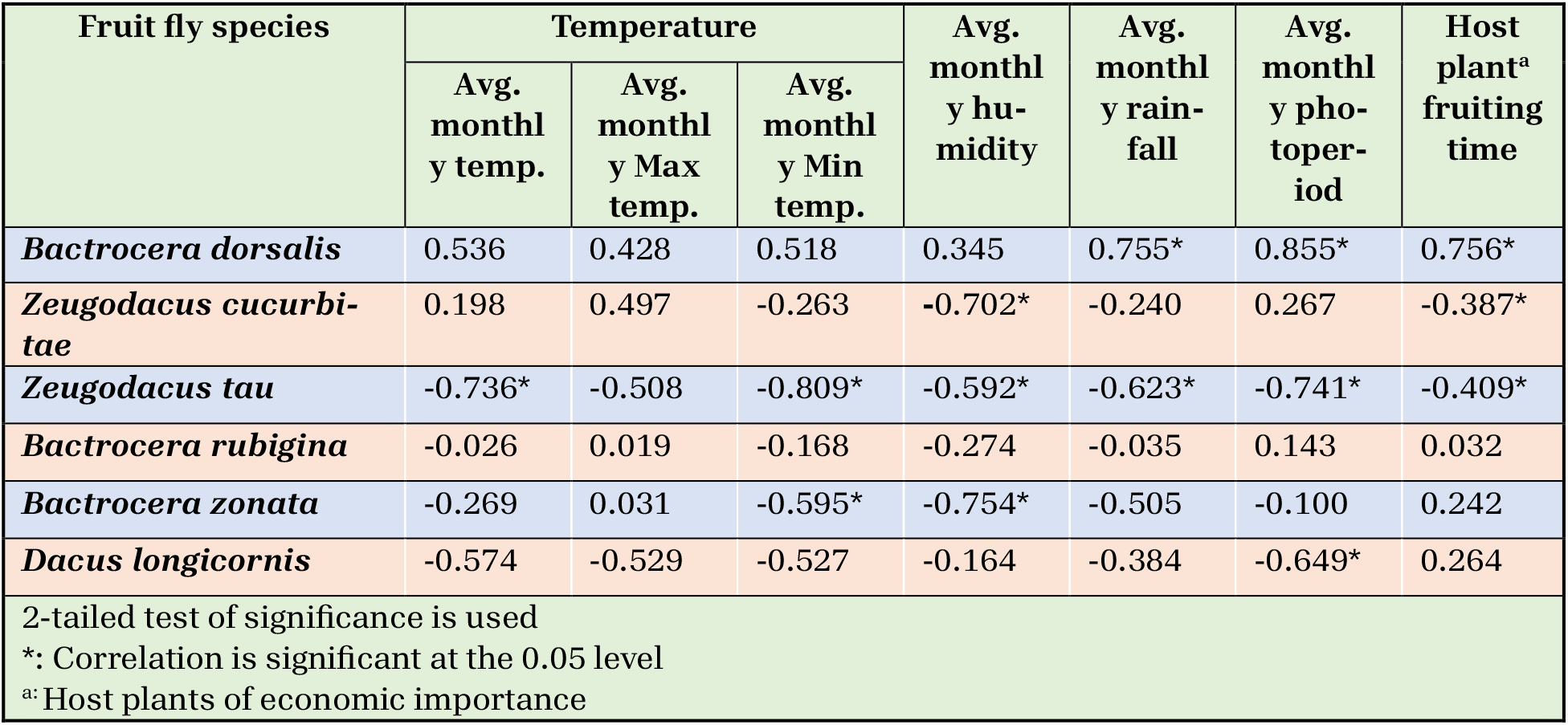
Pearson’s correlation coefficient with weather conditions and host plant fruiting time of six most abundant Tephritidae fruit fly.

**Fig. 6:**
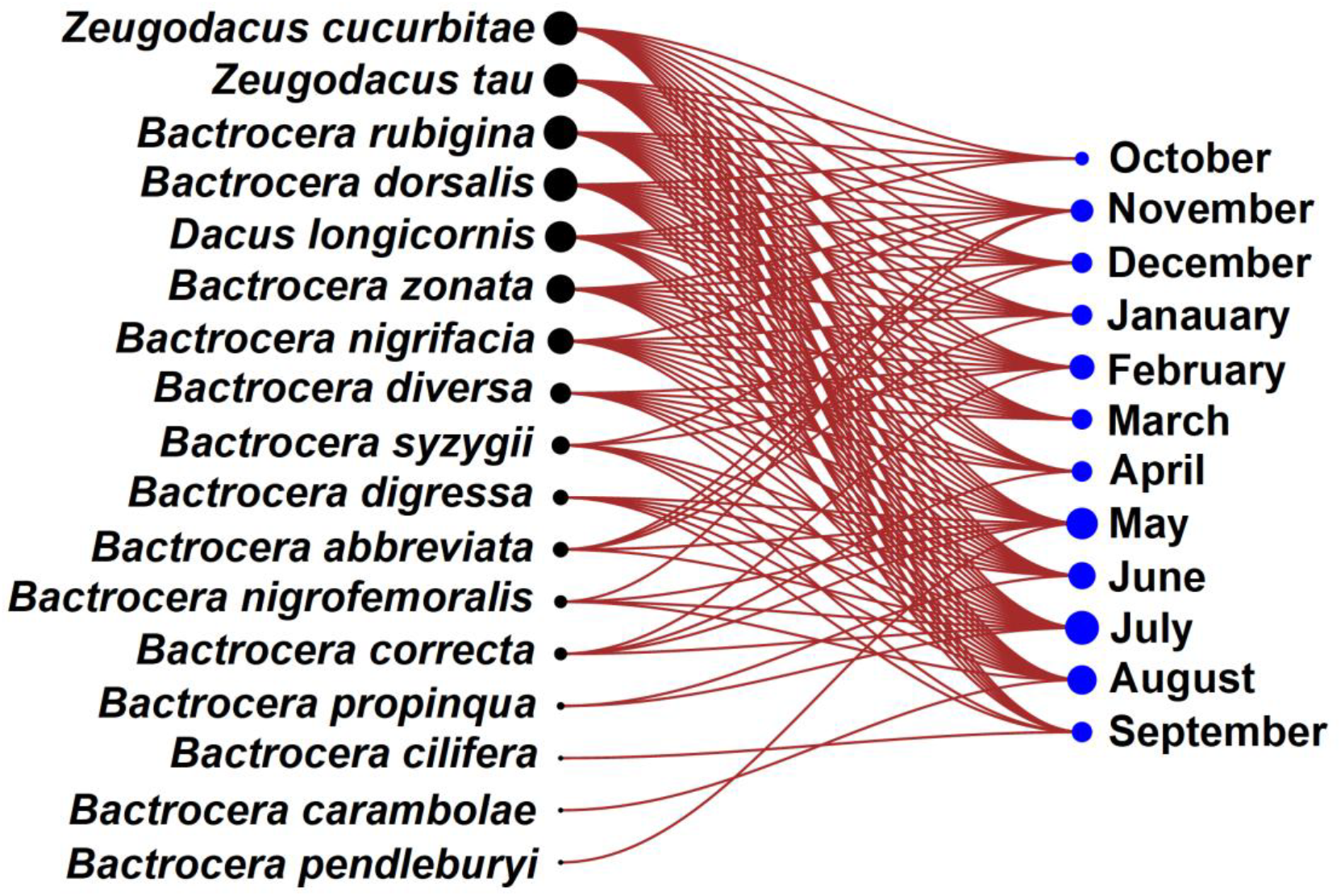
Bipartite linear graphs of the Tephritidae species and their detecting months at AERE campus: black nodes (•) represent the species and blue nodes (•) represent the respective months. The size of black node (•) reflects on the number of species detected (1 to 17) and the blue nodes (•) size reflects the respective number of months concerned (1 to 12).

### 3.4. Correlation of population dynamics with weather conditions and host plant fruiting

We collected meteorological data including temperature, rainfall, relative humidity, and photoperiod (**Fig. 7**) from the local weather station of Bangladesh Meteorological Department, Dhaka, Bangladesh. Fruiting times of economically important host plants in this AERE area during our survey are shown in **Fig. 8**.

**Fig. 7:**
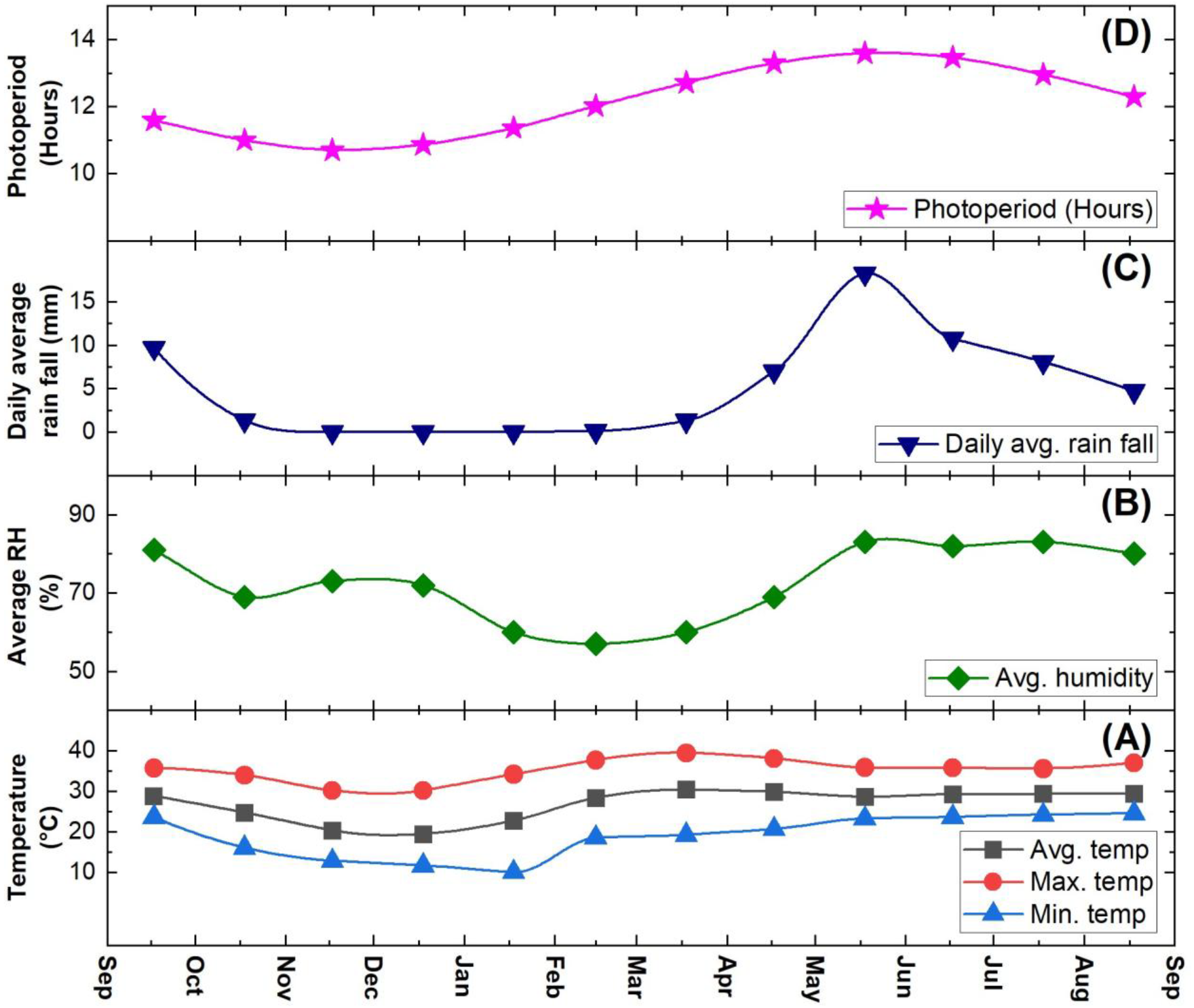
Climatic factors (A) temperature, (B) average relative humidity, (C) daily average rainfall and (D) photoperiod of survey area (AERE) during survey.

**Fig. 8:**
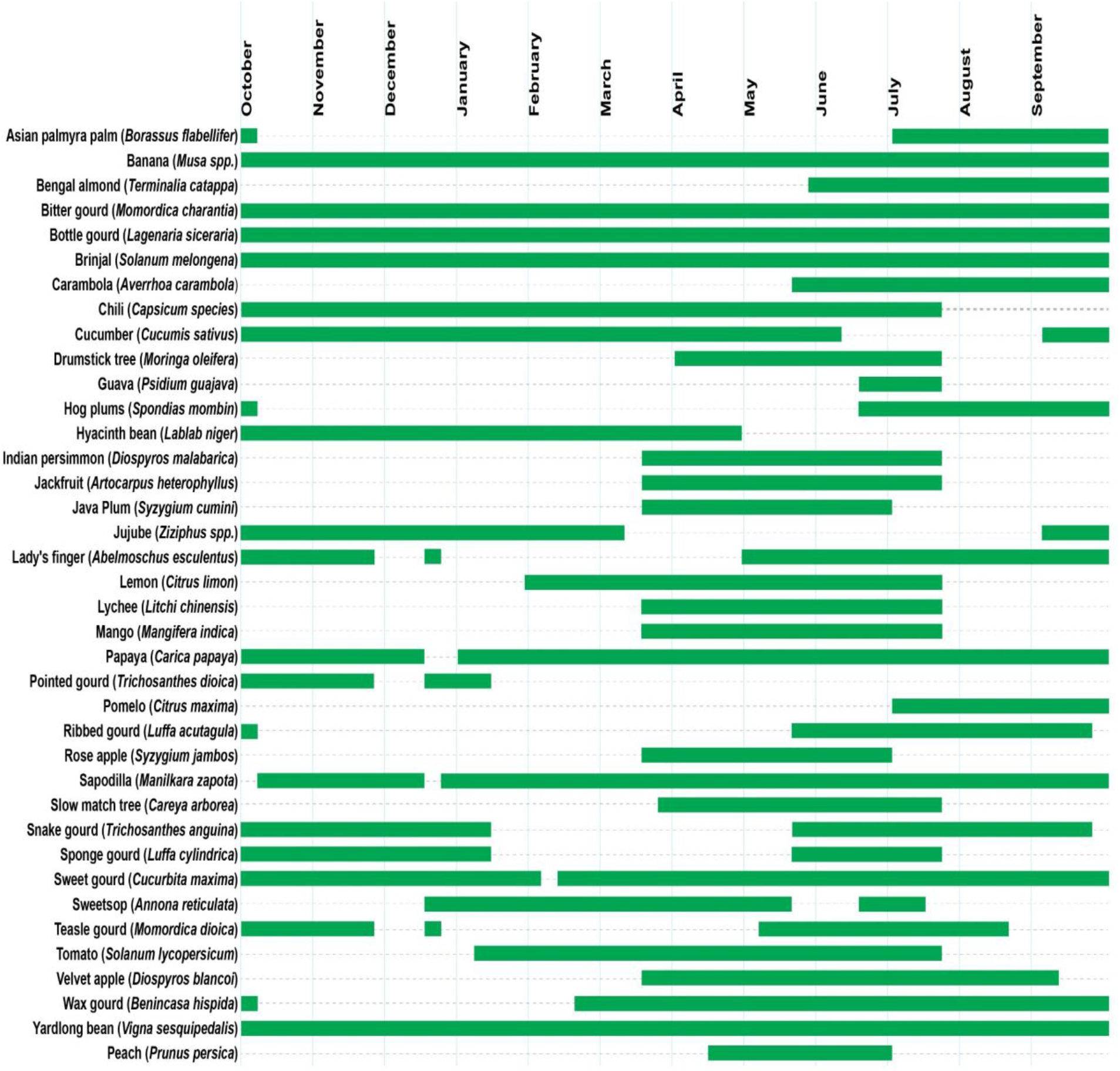
The annual fruiting period (from October to September) for thirty-eight commercially important host plants of Tephritidae fruit flies at AERE area.

The Pearson’s correlation coefficient values in **Table 1** illustrate the relationship between the FTD of the six most abundant Tephritidae fruit fly species and key environmental factors (temperature, rainfall, humidity, photoperiod) as well as fruiting time of host plants (**Fig. 8**) of economic importance. These values provide insight into how changes in these variables may affect the population dynamics of these fruit fly species.

The detailed year-round population dynamics of six Tephritidae fruit fly species have been shown in **Fig. 9** to **Fig. 14**, revealing population trends and possibility of the existence of subgroups.

**Fig. 9:**
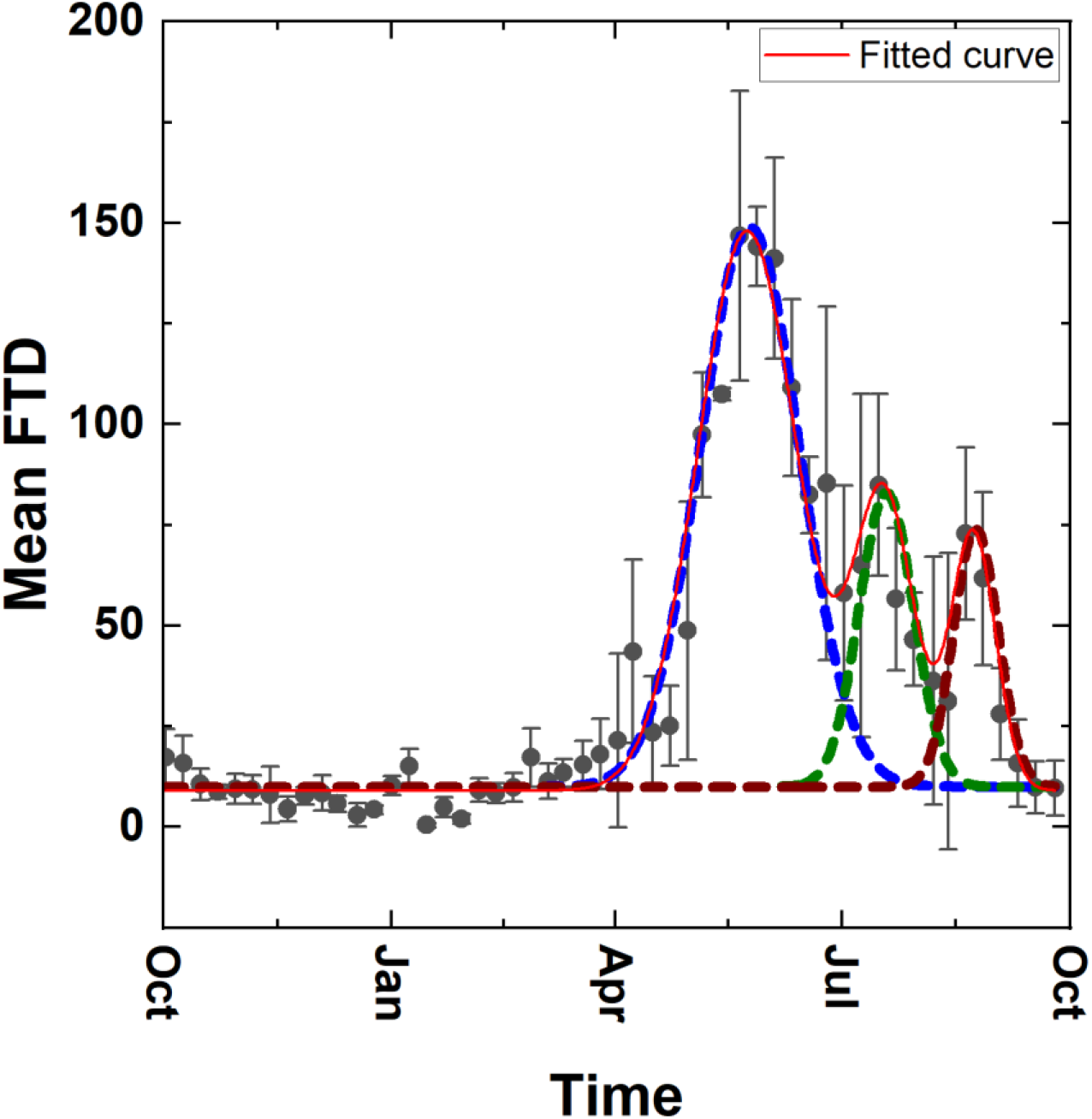
Year-round population dynamics of *Bactrocera dorsalis*. The red color represents the fitted curve obtained from a linear combination of three overlapping Gaussian population distributions. The first population peak appeared on 29^th^ May (represented by dotted blue color Gaussian lineshape). The second population peak was on 21^st^ July (represented by dotted green color Gaussian lineshape) and the third peak was on 28^th^ August (represented by dotted wine color Gaussian lineshape).

#### 3.4.1. Bactrocera dorsalis

*Bactrocera dorsalis* (Hendel) is a pest that reproduces multiple times a year (Jaffar *et al*., 2023) and has a wide range of hosts. This fruit fly has been reported to affect over 250 varieties of fruits and vegetables, causing significant damage to crop production worldwide. (Clarke *et al*., 2005; Qin *et al*., 2018). In addition, this species is able to survive in various weather conditions of the subtropical monsoon climate throughout the year. The contribution of all these factors established it as the predominant pest species within our survey area. **Fig. 9** shows the yearlong survey with the fitted curve obtained from a linear combination of three overlapping Gaussian population distribution. Our Gaussian mixture model suggests that the population dynamics possesses three subgroups over the year along with three distinct growth peaks of *B. dorsalis* population (**Fig. 9**). According to a previous study, *B. dorsalis* is multivoltine, completing several overlapping generations (6.03–8.28) annually (Choudhary *et al*., 2019). Therefore, the presence of these three subgroups suggests the clustering of consecutive overlapped generations of *B. dorsalis* in this survey area (Barclay et al., 2016).

During the trapping period, the counts of the average monthly captured *B. dorsalis* were at their lowest in December 2020 and January 2021, with average monthly FTD around 5.67 and 6.91 respectively. However, the population of this pest species began to increase in March (14.93 average monthly FTD). The average monthly FTD sharply increased to 110.83 in May and 107.19 in June. Afterward, the population of *B. dorsalis* began to decline, and this trend continued up to September 2021. The *B. dorsalis* population consistently remained low from October to March, with an average monthly FTD always below 14.93. The period of low population lasted for about six months.

In this study, the abundance of male *B. dorsalis* and the fruiting stage of host plants (**Fig. 7**) were observed primarily during the wet seasons of the year. Pearson’s correlation analysis (**Table 1**) revealed a significant correlation (p < 0.05) of the population of *B. dorsalis* with the availability of host plant fruiting time, rainfall as well as photoperiod. Similar findings were also reported by Chen *et al*. (Chen *et al*., 2006) and Tan and Serit (Tan & Serit, 1994).

According to Ye and Liu, rainfall affects the population of *B. dorsalis* in two ways (Ye & Liu, 2007). In dry conditions, only 11.67% of the third instar larvae survive and develop to adulthood (Ren Lu *et al*., 2007). On the other hand, adequate rainfall increases soil moisture and reduces mortality rates of mature larvae and facilitates new adult emergence. However, excess precipitation can result in high soil moisture levels, leading to saturated moist conditions. In the saturated moist conditions, none of the larvae and pupae can survive (Ren Lu *et al*., 2007).

In addition, another possible factors for this temporal population dynamics could be attributed to the seasonal variations in reproductive activity of this species as mentioned by Clarke *et al*. (Clarke *et al*., 2022). Moreover, this population dynamics could also be influenced by the trade-off between reproduction and stress-related traits (Chen *et al*., 2017).

#### 3.4.2. Zeugodacus cucurbitae

*Zeugodacus cucurbitae* (Coquillett) is a significant agricultural pest that infests a wide variety of host plants. It has been reported that the *Z. cucurbitae* can infest 125 different host plants (Ahn *et al*., 2022). This species is particularly well adapted to warm climates and can survive in temperatures ranging from 13.9°C to 35.2°C (Ahn *et al*., 2022; Mkiga & Mwatawala, 2015). The remarkable adaptation ability with temperature and ability to infest a wide range of host plants established them as the second most prevalent pest species in our survey area. It was observed that cucurbit vegetables were its primary host in the survey area. **Fig. 10** shows the yearlong survey along with the fitted curve obtained from a linear combination of two overlapping Gaussian population distributions. The green and blue color Gaussian lineshapes have very little overlap around the last half of March

**Fig. 10:**
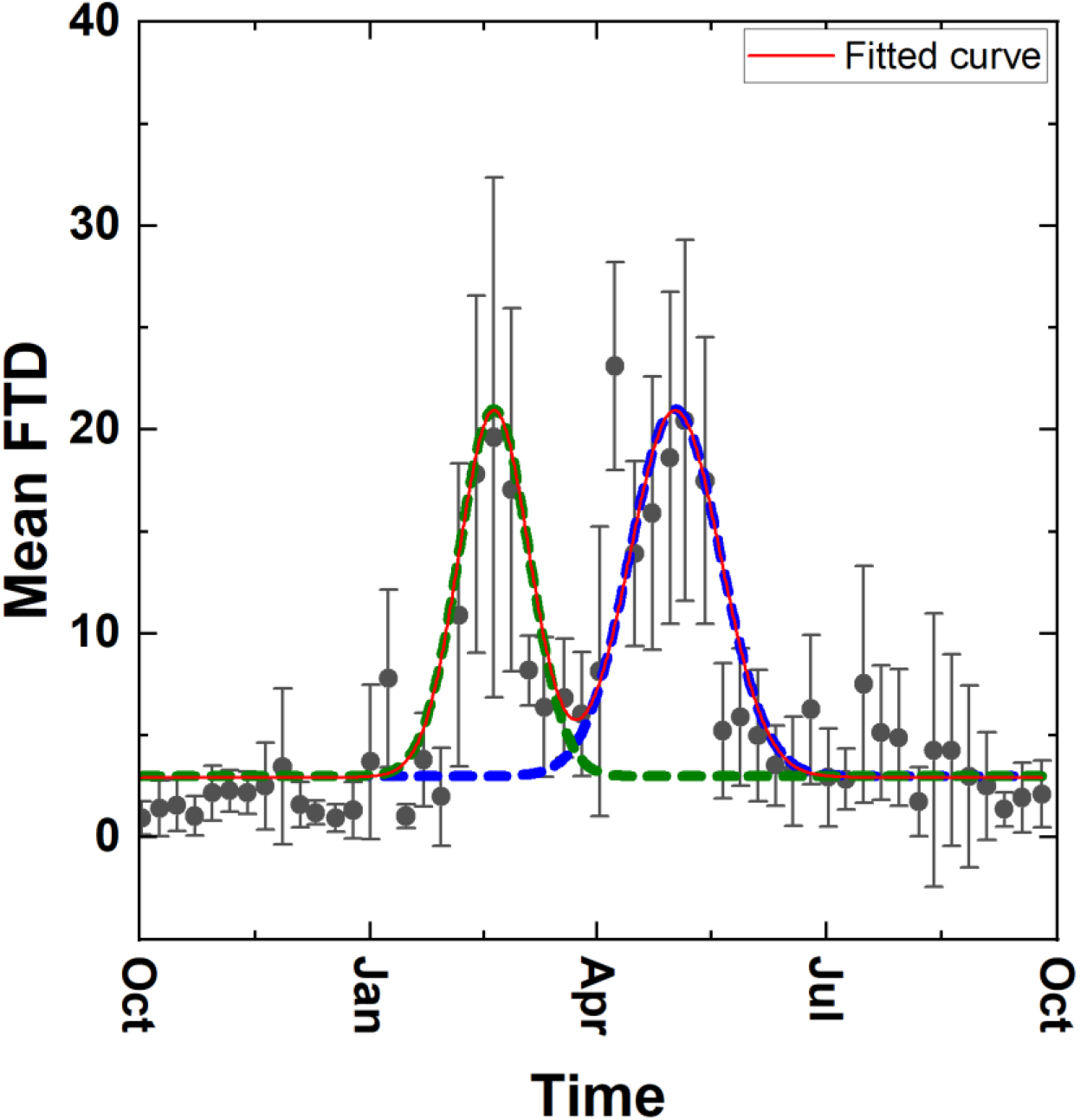
Year-round population dynamics of *Zeugodacus cucurbitae*. The red color represents the fitted curve obtained from a linear combination of two overlapping Gaussian population distributions. The first population peak appeared on the 24^th^ of February (represented by dotted green color Gaussian lineshape). The second population peak appeared on the 7^th^ of May (represented by dotted blue color Gaussian lineshape). Between the two population peaks, the lowest population was observed in the last week of March.

Our Gaussian mixture model suggests that the population dynamics of *Z. cucurbitae* possesses two subgroups over the year along with two distinct growth peaks (**Fig. 10**). According to Dhillon *et al*. there are possibility of many generations (8 to 10) of *Z. cucurbitae* in a year (Dhillon et al., 2005). Therefore, the two distinct growth peaks obtained from the Gaussian mixture model suggest an accumulation of more than one generation for each peak in the population dynamics of the year-long survey (Barclay et al., 2016).

We observed that the population of this pest remained low from October to December, with a range of average monthly FTD value from 1.34 to 2.51. The population reached the lowest point in October 2020, declining to only 1.34 average monthly FTD. However, from January 2021, the population began to increase, reaching 3.84 average monthly FTD value (**Fig. 10**). The first peak was observed in February, with a population of 14.73 average monthly FTD. In March, the population slightly decreased to 7.87 average monthly FTD. The population decline may be a result of the average maximum temperature reaching 37.7°C, which is detrimental for their breeding and survival (Ahn *et al*., 2022). The highest population of *Z. cucurbitae* was recorded in April 2021, with an average monthly FTD of 15.19. However, this data point (**Fig. 10**) does not follow our fitted curve. According to the fitted curve obtained from our Gaussian mixture model, we observed the second population peak appeared on the 7^th^ of May. We observed that during the drier and colder months of the year, the population of this species was significantly lower due to their lower survival rate in low temperatures (Mkiga & Mwatawala, 2015).

Pearson correlation analysis (**Table 1**) also shows that the population density positively correlated with temperature and photoperiod. In addition, Pearson correlation analysis (**Table 1**) revealed a significant negative correlation (p < 0.05) between the *Z. cucurbitae* population and average humidity, as well as host plant fruiting time. The negative correlation with the number of host plant species at the fruiting stage may be due to a decrease in the quantity of available fruit for breeding, even though there is an increase in the number of host plant species at fruiting stage.

#### 3.4.3. Zeugodacus tau

In our study, *Zeugodacus tau* (Walker) was identified as the third most abundant species in terms of population size compared to other fruit fly species. Still, *Z. tau* remains the second most serious fruit fly pest affecting cucurbits in this survey area. In our survey, the highest number of *Z. tau* flies per trap was recorded in February, with an average of 6.65 monthly average FTD. Thereafter, the average population of this pest declined significantly from April onwards and reached its lowest point in September. Over the span of these six months, the monthly average capture range fluctuated from 1.34 to 0.34 FTD. Pearson correlation analysis (**Table 1**) shows a significant (p < 0.05) negative correlation of adult male capture rate with monthly average minimal temperature, average rainfall, humidity, and photoperiod. The higher population count during the cooler month of February could be due to its ability to survive under adverse temperatures spanning −5 to 0 °C and 39 to 42 °C (Liu *et al*., 2022).

This findings is consistent with previous research conducted by Li *et al*. (Li *et al*., 2020). In addition, *Z. tau* exhibited a significant (p < 0.05) negative correlation (**Table 1**) with the fruiting time of its host plants, which was similar to that of *Z. cucurbitae*. As *Z. cucurbitae* and *Z. tau* are also closely related and often share similar ecological niches, they have similar host range and may compete for the same resources for survival. It was reported that *Z. cucurbitae* can outperform *Z. tau* in their shared host environment during interspecific competition, particularly at highdensity as well as low and high temperatures (Shen *et al*., 2014). This is one of the main reasons for the higher population of *Z. cucurbitae* compared to *Z. tau* in our survey area (**Fig. 3**). Because of their very low count rate, there is a high level of uncertainty associated with the mean FTD. Therefore, the fitted curve with data in **Fig. 11** only represents an overall roughly Gaussian like distribution of population with high uncertainty. However, this fitted curve in **Fig. 11** is useful to detect the broad single population peak with seasonal fluctuations in population.

**Fig. 11:**
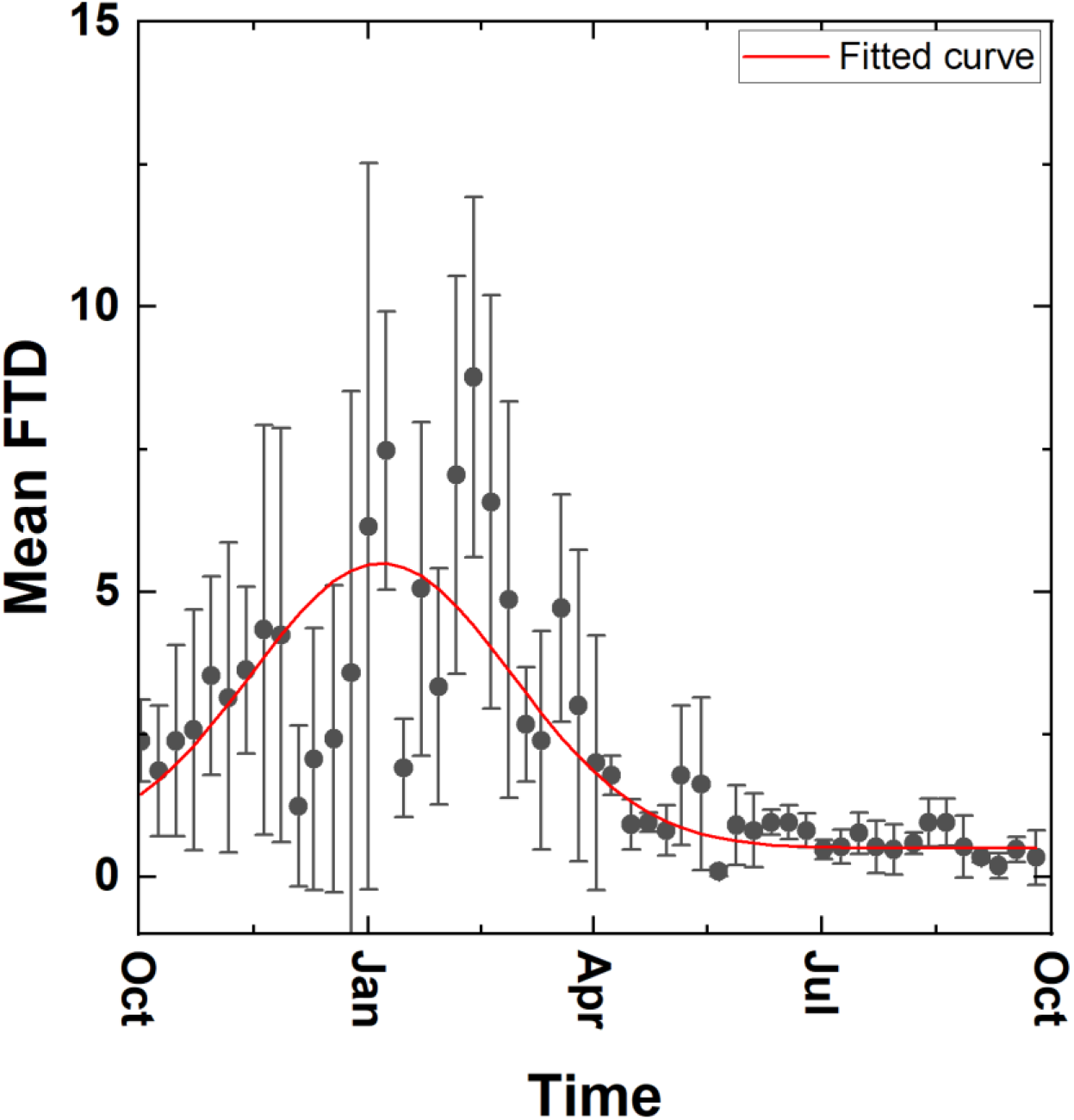
Year-round population dynamics of *Zeugodacus tau*. The red color line represents fitted curve of population density following a roughly Gaussian distribution, reaching the maximum population density on 11^th^ of January.

According to Li *et al*., this species has three to four generations each year, with significant overlap between generations (Li et al., 2020). Therefore, the single peak in population distribution in **Fig. 11** indicates the overlapping of several generations between October and April.

#### 3.4.4. Bactrocera rubigina

*Bactrocera rubigina* (Wang & Zhao) is not classified as a pest species, as there is no evidence of significant economic impacts associated with this species (Drew *et al*., 2007). This species was detected all year round, possibly due to their consistent breeding behavior (**Fig. 12**). The consistent presence of this species in our survey was likely due to the overlapping generations and the presence of all life stages at any given time.

**Fig. 12:**
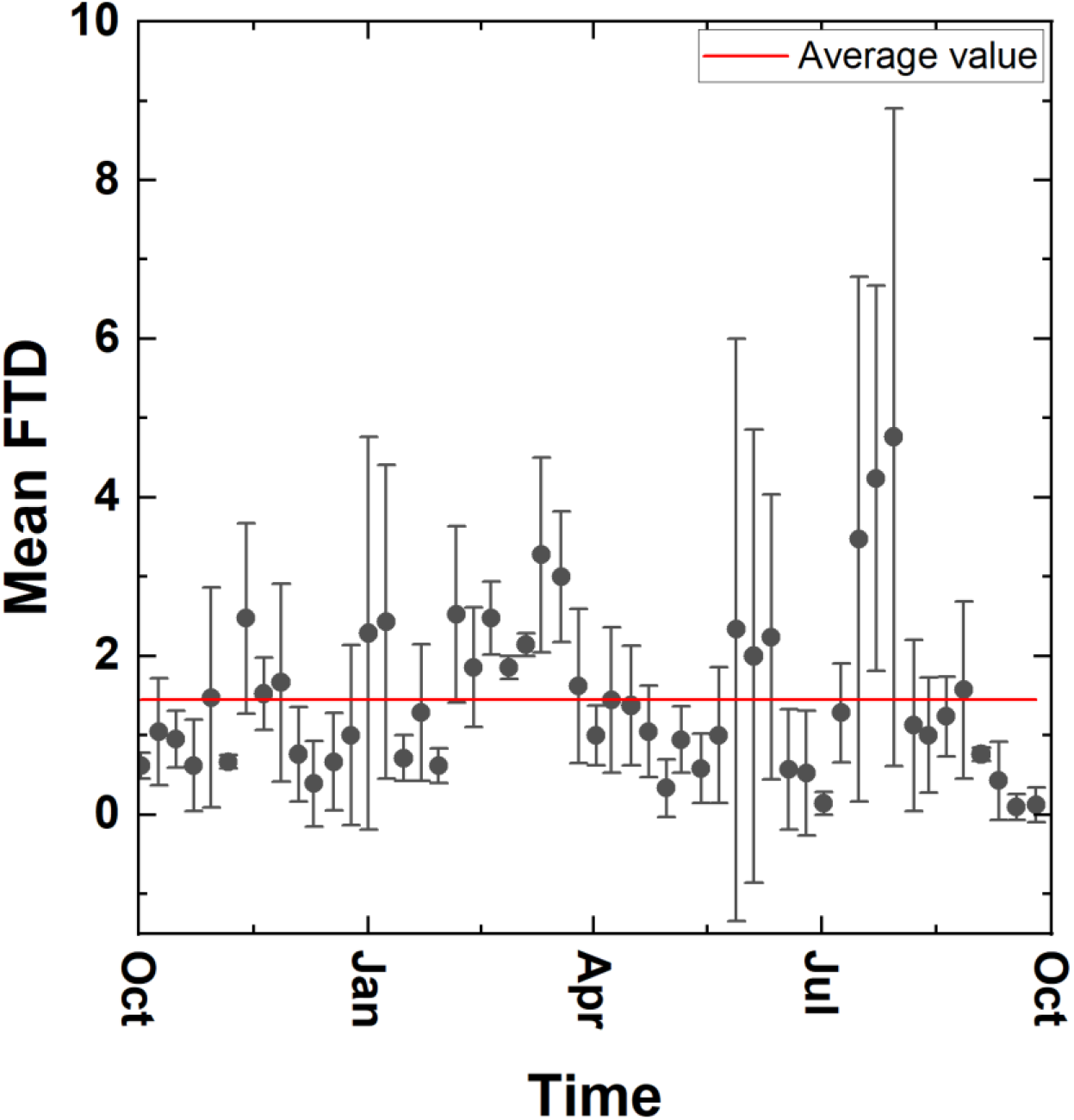
Year-round population dynamics of *Bactrocera rubigina*. The red color represents the average value of the population over the year. No distinct peak of population density was observed as the captured mean FTD was very low.

It is likely that this species maintains a stable population throughout the year because of their reproductive patterns. The red line in **Fig. 12** represents the average FTD in our study period. We observed an average monthly FTD value in the range from 0.39 to 2.53. The highest captured FTD was between 20^th^ July to 04^th^ August, whereas the lowest value was observed in September (**Fig. 12**). However, there is high uncertainty associated with the mean FTD value because the captured mean FTD was extremely low.

Consequently, no distinct peak in population density was observed. Pearson correlation analysis (**Table 1**) also does not show any significant correlation of *B. rubigina* population dynamics with weather conditions or host plant fruiting time. However, despite being detected throughout the year, its presence does not pose a threat to crops or agricultural production.

The fluctuating FTD values indicate varying levels of activity of this species. Overall, the species is not of major concern in terms of pest management. According to available reports, the sage-leaved alangium (*Alangium salviifolium*) is identified as a host plant that bears fruits during the months of March to May (Guang-qin *et al*., 1993). However, there is a possibility of undiscovered host plants that still need to be identified. Since very limited details are available about their host species (Drew *et al*., 2007), more comprehensive research is required to uncover the hidden unknown hosts and the impact of this species on agriculture

#### 3.4.5. Bactrocera zonata

Throughout the one-year trapping period, *Bactrocera zonata* (Saunders) was detected from November to August, with fluctuating average monthly FTD values ranging from 0.01 to 0.14. However, no flies of this species were detected in October and September. Comparatively higher abundance of this species was observed from January to July (**Fig. 13**). The FTD value peaked at 0.14 in April, indicating a prominent presence of *B. zonata* during that month. Due to low captured rate and high uncertainty (represented by the error bars) associated with the mean FTD data, it was not possible to find any pattern of *B. zonata* population dynamics in the study period. However, the fitted curve to data using our Gaussian distribution model suggested a single broad peak of population density from January to July, reaching its highest point around April. (**Fig. 13**).

**Fig. 13:**
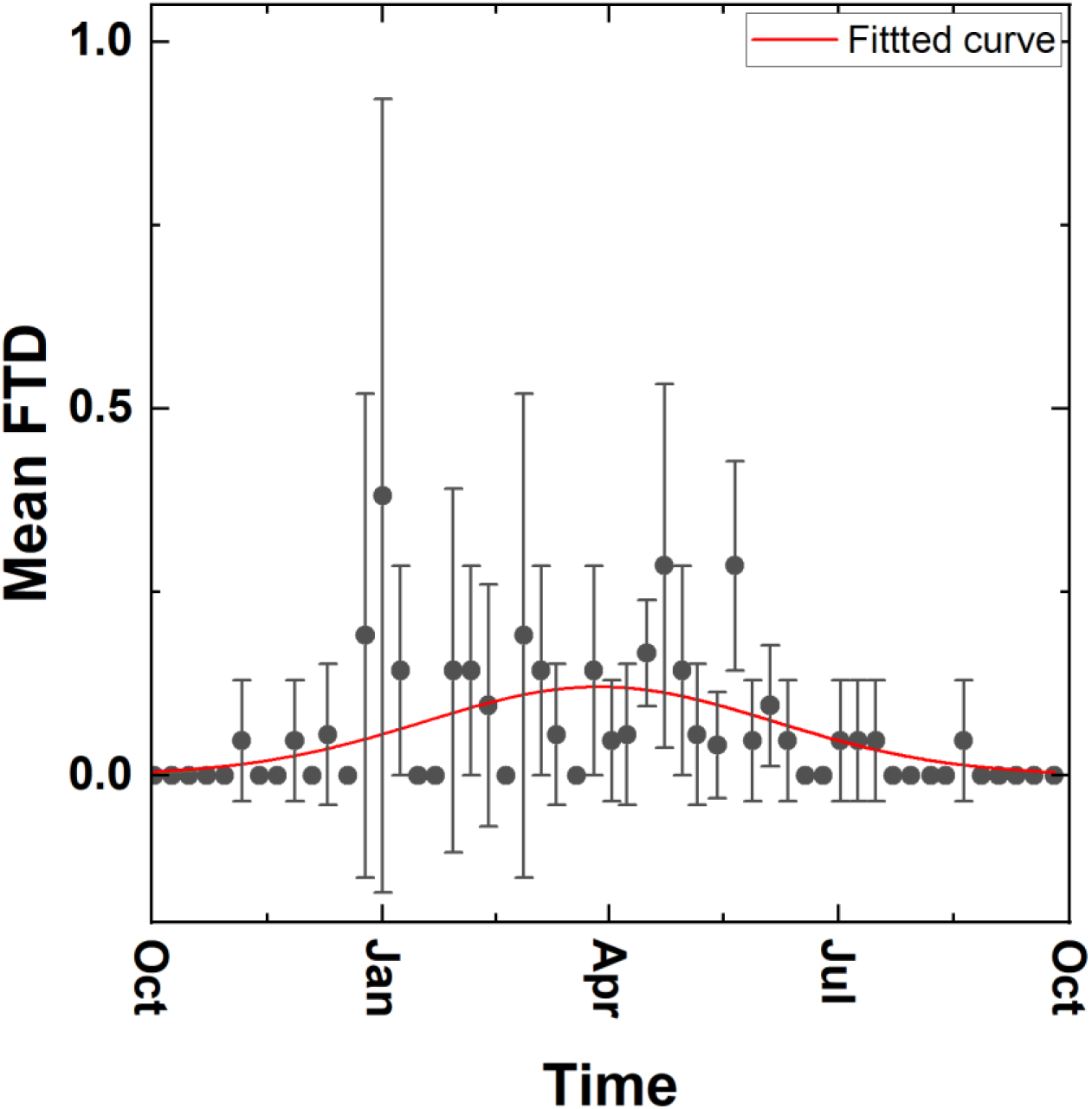
Year-round population dynamics of *Bactrocera zonata*. The red color curve represents the fitted curve to data of a roughly Gaussian population distribution with a broad peak. Due to low captured rate and high uncertainty associated with mean the FTD data, the fitted curve only suggests a possible maximum population density around April.

In our study, we discovered that *B. zonata* and *B. dorsalis* had significantly different FTD values. Specifically, the annual average FTD value for *B. zonata* was 0.06, while for *B. dorsalis* the annual average FTD value was 36.33. In a study by Moquet *et al*. in 2021, the authors suggested that these two pest species may struggle to coexist in the same host range.

According to their research, the invasion of *B. dorsalis* reduced the diversity and richness of host range of *B. zonata* by 50% (Moquet *et al*., 2021). In our study area, both *B. zonata* and *B. dorsalis* share the same host plants. The small population size of *B. zonata* may be due to competition with the highly competitive pest *B. dorsalis* for resources on these shared host plants.

Pearson correlation analysis (**Table 1**) revealed a significant negative correlation (p < 0.05) of the population of *B. zonata* with average monthly humidity and average monthly minimum temperature. A study by Duyck *et al*. revealed that ovarian maturation of females only occurs within a narrow temperature range of 25°C to 30°C (Duyck *et al*., 2004). This suggests that average monthly minimum temperature while going outside of this range could have an impact on their reproductive success. In our case, monthly minimum temperature falling outside of this range may have affected their reproductive success. However, according to a recent study, *B. zonata* has a potential to expand its geographical range to new regions with suitable climates across the world (Bayoumy *et al*., 2021; Ullah *et al*., 2023)

#### 3.4.6. Dacus longicornis

*Dacus longicornis* (Wiedemann) is a pest of many cucurbit crops. In our study area, we observed a very low capture rate of *D. longicornis* in the fruit fly traps, with average monthly capture rates (FTD) ranging from 0.006 to 0.21. However, we observed comparatively higher average monthly FTD from October to March (**Fig. 14**) with a fluctuating trend. The highest population density was recorded on December 22^nd^. In the course of this survey, it is noteworthy that no *D. longicornis* was detected in the month of April. One potential explanation for their unusually small population size could be the intense competition they face from *Z. cucurbitae* and *Z. tau*. This competition arises because all three species share the same host range, resulting in limited resources for each species to thrive and reproduce. Among the abiotic factors, the Pearson correlation analysis (**Table 1**) revealed a significant negative correlation (p < 0.05) between the population of *D. longicornis* and the average monthly photoperiod. Also, Pearson correlation analysis shows a weak positive correlation with the fruiting time of host plants

**Fig. 14:**
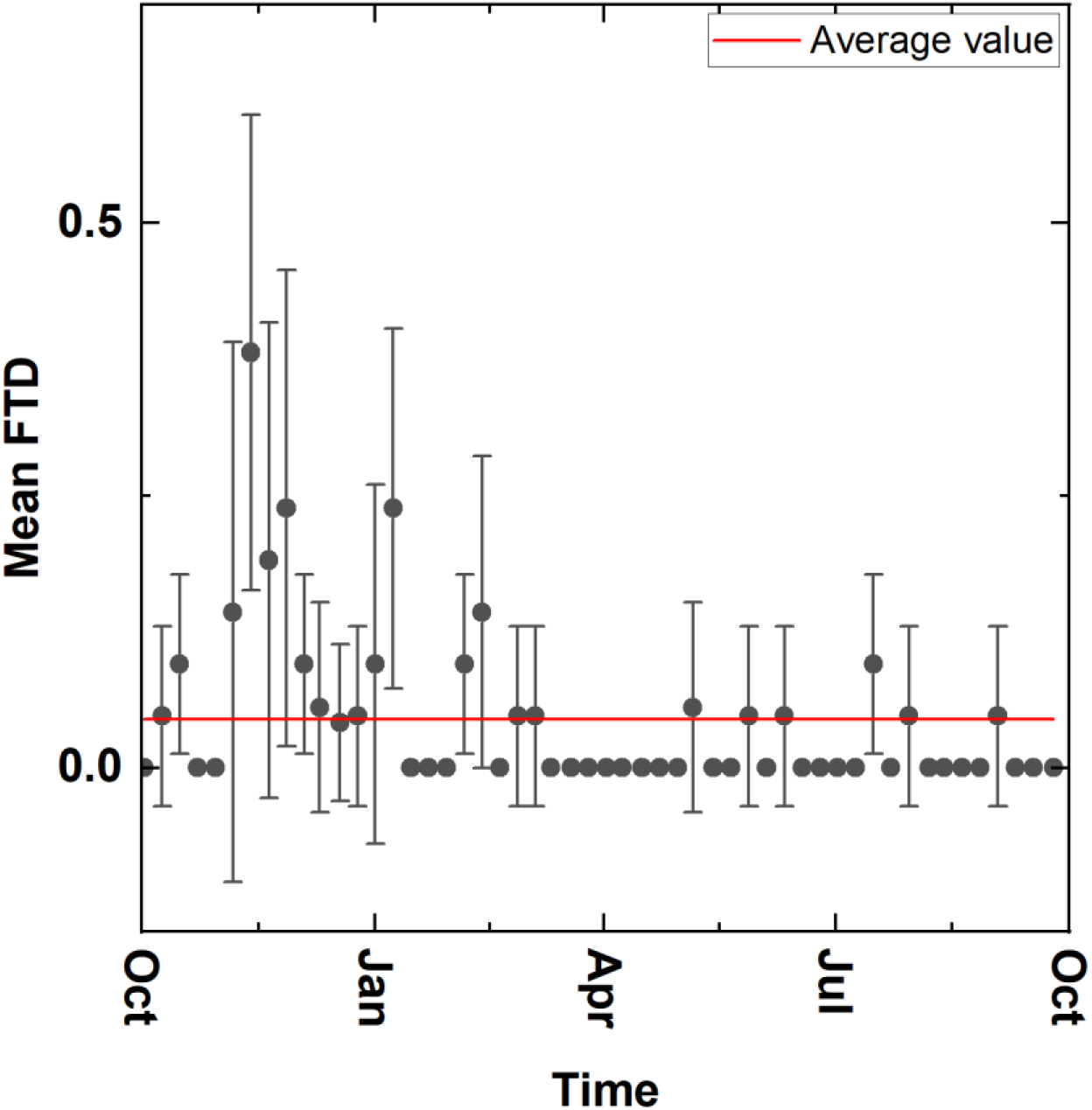
Year-round population dynamics of *Dacus longicornis*. The red color represents the average value of the population over the year.

## 4. Conclusion

Our year-round investigation within the semirural environment of Atomic Energy Research Establishment, Dhaka, Bangladesh presents a detailed statistical analysis of Tephritidae fruit flies population dynamics, species richness and diversity as well as comparison of detection range of three widely used male lures parapheromone traps. A total of seventeen Tephritidae fruit fly species were detected during this survey. Out of the seventeen species, only six Tephritidae fruit fly species (*B. dorsalis, Z. cucurbitae, Z. tau, B. rubigina, B. zonata*, and *D. longicornis*) were found to be more abundant throughout the year-long survey. This survey detected *B. carambola* for the first time in this area, indicating the spread of this species towards north-west region of Bangladesh. Top two abundant pest fruit fly species, namely *B. dorsalis* and *Z. cucurbitae*, have been consistently observed at infestation levels (>1 FTD) throughout the year, while FTD of *Z. tau* reaches infestation levels (>1 FTD) from October to May. Other detected pest fruit flies average monthly FTD values were either at suppression level (1.0 to 0.1 FTD) or eradication level (<0.1 FTD) (IAEA, 2003). Our study suggests that it is crucial to implement necessary control measures to suppress the abundant pest fruit fly species namely *B. dorsalis, Z. cucurbitae*, and *Z. tau*, comprising 96.8% of the total count. After reducing pest populations to the required threshold level, the Sterile Insect Technique (SIT) and other control measures could be implemented to eradicate the remaining fruit fly pests. However, it is crucial to note that non-pest fruit fly populations should also be carefully monitored, as they can reach infestation levels for an extended period of time. For instance, *B. rubigina* was found to be above 1 FTD for approximately eight months

In addition, our statistical analysis using Pearson’s correlation coefficient confirms that the population dynamics of the most abundant six fruit fly species detected in the survey area were affected by the weather conditions of their habitat as well as the availability of suitable fruiting plants for their breeding and survival. For the first time, our mathematical model based on the Gaussian mixture model indicated the presence of the overlapped subgroups within the population of the top two most abundant species (*B. dorsalis* and *Z. cucurbitae*) over the yearlong period in this area. This mathematical modelbased approach can be applied to other insect species with sufficient counts to reveal the hidden nature of their population dynamics. This data-driven mathematical model-based approach would allow for taking informed decision regarding an efficient integrated pest management of pestiferous fruit fly species, minimizing the impact of fruit fly infestations.

## CRediT authorship contribution statement

**Mahfuza Momen:** Conceptualization, Data curation, Formal Analysis, Funding acquisition, Investigation, Methodology, Project administration, Resources, Software, Supervision, Validation, Visualization, Writing original draft, Writing review & editing. **Md. Aftab Hossain:** Funding acquisition, Project administration, Resources, Writing review & editing. **Kajla Sheheli:** Funding acquisition, Writing review & editing. **Md. Forhad Hossain:** Data curation, Investigation. **Md. Abdul Bari:** Writing-review & editing.

## Funding

This study was supported by the Bangladesh Atomic Energy Commission (BAEC) in Bangladesh under project IFRB-IBD-1/2002.

## Declaration of competing interest

The authors do not have any financial or non-financial interests to disclose in relation to this article. The authors have no competing interests to declare that are relevant to the content of this article.

## Data availability

The experimental data will be provided upon reasonable request.

## Notes

### Competing Interest Statement

The authors have declared no competing interest.

### Summary of Updates

Updated map (Figure 1) with copyright attribution, and corrected typos

## References

Agarwal, M., & Sueyoshi, M. (2005). Catalogue of Indian fruit flies (Diptera: Tephritidae). Oriental Insects - ORIENT INSECT, 39, 371–433. 10.1080/00305316.2005.10417450

Ahn, J. J., Choi, K. S., & Huang, Y.-B. (2022). Thermal effects on the development of Zeugodacus cucurbitae (Coquillett) (Diptera: Tephritidae) and model validation. Phytoparasitica, 50(3), 601–616. 10.1007/s12600-022-00985-5

Barclay, H. J., Enkerlin, W., Manoukis, N. C., & Flores, J. R. (2016). Guidelines for the use of mathematics in operational area-wide integrated pest management programmes using the sterile insect technique with a special focus on Tephritid fruit flies. FAO/IAEA. https://www.iaea.org/sites/default/files/21/06/nafa-ipc-manual-tephritid-fruit-flies-manual.pdf

Bayoumy, M. H., Michaud, J. P., Badr, F. A. A., & Ghanim, N. M. (2021). Validation of degree-day models for predicting the emergence of two fruit flies (Diptera: Tephritidae) in northeast Egypt. Insect Sci, 28(1), 153–164. 10.1111/1744-7917.12750

Chao, A. (1984). Nonparametric Estimation of the Number of Classes in a Population. Scandinavian Journal of Statistics, 11(4), 265–270. http://www.jstor.org/stable/4615964

Chao, A., & Shen, T.-J. (2003). Nonparametric estimation of Shannon’s index of diversity when there are unseen species in sample. Environmental and Ecological Statistics, 10(4), 429–443. 10.1023/A:1026096204727

Chen, E.-H., Hou, Q.-L., Wei, D.-D., Jiang, H.-B., & Wang, J.-J. (2017). Phenotypes, antioxidant responses, and gene expression changes accompanying a sugar-only diet in Bactrocera dorsalis (Hendel) (Diptera: Tephritidae). BMC Evolutionary Biology, 17(1). 10.1186/s12862-017-1045-5

Chen, P., Ye, H., & Liu, J. (2006). Population dynamics of Bactrocera dorsalis (Diptera: Tephritidae) and analysis of the factors influencing the population in Ruili, Yunnan Province, China. Acta Ecologica Sinica, 26(9), 2801–2808. 10.1016/S1872-2032(06)60044-9

Chiu, C. H., Wang, Y. T., Walther, B. A., & Chao, A. (2014). An improved nonparametric lower bound of species richness via a modified good-turing frequency formula. Biometrics, 70(3), 671–682. 10.1111/biom.12200

Choudhary, J. S., Mali, S. S., Mukherjee, D., Kumari, A., Moanaro, L., Rao, M. S., Das, B., Singh, A. K., & Bhatt, B. P. (2019). Spatio-temporal temperature variations in MarkSim multimodel data and their impact on voltinism of fruit fly, Bactrocera species on mango. Scientific Reports, 9(1), 9708. 10.1038/s41598-019-45801-z

Clarke, A. R., Armstrong, K. F., Carmichael, A. E., Milne, J. R., Raghu, S., Roderick, G. K., & Yeates, D. K. (2005). INVASIVE PHYTOPHAGOUS PESTS ARISING THROUGH A RECENT TROPICAL EVOLUTIONARY RADIATION: The Bactrocera dorsalis Complex of Fruit Flies. Annual Review of Entomology, 50(Volume 50, 2005), 293–319. 10.1146/annurev.ento.50.071803.130428

Clarke, A. R., Leach, P., & Measham, P. F. (2022). The Fallacy of Year-Round Breeding in Polyphagous Tropical Fruit Flies (Diptera: Tephritidae): Evidence for a Seasonal Reproductive Arrestment in Bactrocera Species. Insects, 13(10), 882. 10.3390/insects13100882

Colwell, R. K., Mao, C. X., & Chang, J. (2004). INTERPOLATING, EXTRAPOLATING, AND COMPARING INCIDENCE-BASED SPECIES ACCUMULATION CURVES. Ecology, 85(10), 2717–2727. 10.1890/03-0557

Dhillon, M. K., Singh, R., Naresh, J. S., & Sharma, H. C. (2005). The melon fruit fly, Bactrocera cucurbitae: A review of its biology and management. Journal of Insect Science, 5(1). 10.1093/jis/5.1.40

Doorenweerd, C., Leblanc, L., Norrbom, A. L., Jose, M. S., & Rubinoff, D. (2018). A global checklist of the 932 fruit fly species in the tribe Dacini (Diptera, Tephritidae). ZooKeys(730), 19-56. 10.3897/zookeys.730.21786

Drew, R., Romig, M., & Dorji, C. (2007). Records of Dacine Fruit Flies and New Species of Dacus (Diptera: Tephritidae) in Bhutan. Raffles Bulletin of Zoology, 55(1). 10.5281/zenodo.5331152

Drew, R. A. I., & Romig, M. C. (2013). Tropical Fruit Flies (Tephritidae Dacinae) of South-East Asia: Indomalaya to North-West Australasia. CABI. 10.1079/9781780640358.0000

Duyck, P.-F., Sterlin, J. F., & Quilici, S. (2004). Survival and development of different life stages of Bactrocera zonata (Diptera: Tephritidae) reared at five constant temperatures compared to other fruit fly species. Bulletin of Entomological Research, 94, 89–93. 10.1079/BER2003285

Guang-qin, Liang, Hancock1, David L., Wei, X., & Fan, L. (1993). NOTES ON THE DACINAE OF SOUTHERN CHINA (DIPTERA: TEPHRITIDAE). Australian Journal of Entomology, 32(2), 137–140. 10.1111/j.1440-6055.1993.tb00561.x

Hossain, M. A., Leblanc, L., Momen, M., Bari, M. A., & Khan, S. A. (2019). Seasonal abundance of economically important fruit flies (Diptera: Tephritidae: Dacinae) in Bangladesh, in relation to abiotic factors and host plants. http://hdl.handle.net/10125/65002

IAEA. (2003). Trapping Guidelines for Area-wide Fruit Fly Programmes. International Atomic Energy Agency. https://www.iaea.org/publications/6916/trapping-guidelines-for-area-wide-fruit-fly-programmes

Jaffar, S., Rizvi, S. A. H., & Lu, Y. (2023). Understanding the Invasion, Ecological Adaptations, and Management Strategies of Bactrocera dorsalis in China: A Review. Horticulturae, 9(9), 1004. 10.3390/horticulturae9091004

Jayakumari, D., Hinde, J., Einbeck, J., & Moral, R. A. (2023). Tools for Assessing Goodness of Fit of GLMs: Case Studies in Entomology. In R.A. Moral & W. A. C. Godoy (Eds.), Modelling Insect Populations in Agricultural Landscapes (pp. 211–235). Springer International Publishing. 10.1007/978-3-031-43098-5_11

Kapoor, V. C. (1970). Indian Tephritidae with their recorded hosts. Oriental Insects, 4(2), 207–251. 10.1080/00305316.1970.10433957

Kapoor, V. C., Hardy, D. E., Agarwal, M. L., & Grewal, J. S. (1980). Fruit fly (Diptera, Tephritidae) : systematics of the Indian subcontinent. Export India Publications Jullundur.

Lan, B. L., & Chandran, P. (2011). Distribution of animal population fluctuations. Physica A: Statistical Mechanics and its Applications, 390(7), 1289–1294. 10.1016/j.physa.2010.11.015

Leblanc, L., Hossain, M. A., Doorenweerd, C., Khan, S. A., Momen, M., San Jose, M., & Rubinoff, D. (2019). Six years of fruit fly surveys in Bangladesh: a new species, 33 new country records and discovery of the highly invasive Bactrocera carambolae (Diptera, Tephritidae). ZooKeys, 876. 10.3897/zookeys.876.38096

Li, X., Yang, H., Hu, K., & Wang, J. (2020). Temporal dynamics of Bactrocera (Zeugodacus) tau (Diptera: Tephritidae) adults in north Jiangxi, a subtropical area of China revealed by eight years of trapping with cuelure. Journal of Asia-Pacific Entomology, 23(1), 1–6. 10.1016/j.aspen.2019.10.007

Liu, H., Wang, X., Chen, Z., & Lu, Y. (2022). Characterization of Cold and Heat Tolerance of Bactrocera tau (Walker). Insects, 13(4), 329. 10.3390/insects13040329

Manly, B. F. J. (1974). Estimation of stage-specific survival rates and other parameters for insect populations developing through several stages. Oecologia, 15(3), 277–285. 10.1007/BF00345183

Mkiga, A. M., & Mwatawala, M. W. (2015). Developmental Biology of Zeugodacus cucurbitae (Diptera: Tephritidae) in Three Cucurbitaceous Hosts at Different Temperature Regimes. J Insect Sci, 15(1). 10.1093/jisesa/iev141

Moquet, L., Payet, J., Glenac, S., & Delatte, H. (2021). Niche shift of tephritid species after the Oriental fruit fly (Bactrocera dorsalis) invasion in La Réunion. Diversity and Distributions, 27(1), 109–129. 10.1111/ddi.13172

Motulsky, H. (2018). Intuitive biostatistics: a nonmathematical guide to statistical thinking. Oxford University Press, USA.

Onufrieva, K. S., & Onufriev, A. V. (2018). Linear relationship between peak and season-long abundances in insects. PLOS ONE, 13(2), e0193110. 10.1371/journal.pone.0193110

Qin, Y.-j., Krosch, M. N., Schutze, M. K., Zhang, Y., Wang, X.-x., Prabhakar, C. S., Susanto, A., Hee, A. K. W., Ekesi, S., Badji, K., Khan, M., Wu, J.-j., Wang, Q.-l., Yan, G., Zhu, L.-h., Zhao, Z.-h., Liu, L.-j., Clarke, A. R., & Li, Z.-h. (2018). Population structure of a global agricultural invasive pest, Bactrocera dorsalis (Diptera: Tephritidae). Evolutionary Applications, 11(10), 1990–2003. 10.1111/eva.12701

Ren Lu, R. L., Lu YongYue, L. Y., & Zeng Ling, Z. L. (2007). Effect of sand water content on the pupal survival of Bactrocera dorsalis (Hendel). Journal of South China Agricultural University, 28(1), 63–66. 10.7671/j.issn.1001-411X.2007.01.015

Reynolds, D. (2009). Gaussian Mixture Models. In S.Z. Li & A. Jain (Eds.), Encyclopedia of Biometrics (pp. 659–663). Springer US. 10.1007/978-0-387-73003-5_196

Scolari, F., Valerio, F., Benelli, G., Papadopoulos, N. T., & Vaníčková, L. (2021). Tephritid Fruit Fly Semiochemicals: Current Knowledge and Future Perspectives. Insects, 12(5), 408. 10.3390/insects12050408

Senior-White, R. (1922). Notes on Indian Diptera. 1. Diptera from the Khasia Hills. 2. Tabanidae in the collection of the forest zoologist. 3. New species of Diptera from the Indian Region. Mem. Dep. Agric. India Entomol. Ser. (Vol. VII (9)). Thacker, Spink & Company., CALCUTTA.

Shen, K., Hu, J., Wu, B., An, K., Zhang, J., Liu, J., & Zhang, R. (2014). Competitive Interactions between Immature Stages of Bactrocera cucurbitae (Coquillett) and Bactrocera tau (Walker) (Diptera: Tephritidae) under Laboratory Conditions. Neotropical Entomology, 43(4), 335–343. 10.1007/s13744-014-0224-y

Soulsby, R. L., & Thomas, J. A. (2012). Insect population curves: modelling and application to butterfly transect data. Methods in Ecology and Evolution, 3(5), 832–841. 10.1111/j.2041-210X.2012.00227.x

Tan, K.-h., & Nishida, R. (2000). Mutual Reproductive Benefits Between a Wild Orchid, Bulbophyllum patens, and Bactrocera Fruit Flies via a Floral Synomone. Journal of Chemical Ecology, 26(2), 533–546. 10.1023/A:1005477926244

Tan, K.-H., & Serit, M. (1994). Adult Population Dynamics of Bactrocera dorsalis (Diptera: Tephritidae) in Relation to Host Phenology and Weather in Two Villages of Penang Island, Malaysia. Environmental Entomology, 23(2), 267–275. 10.1093/ee/23.2.267

Ullah, F., Zhang, Y., Gul, H., Hafeez, M., Desneux, N., Qin, Y., & Li, Z. (2023). Estimation of the potential geographical distribution of invasive peach fruit fly under climate change by integrated ecological niche models. CABI Agriculture and Bioscience, 4(1), 46. 10.1186/s43170-023-00187-x

Van Os, K. (2022). Modelling the timing of generations of Queensland fruit fly, Bactrocera tryoni [Master of Science, University of Canterbury]. 10.26021/12557

Vargas, R. I., Shelly, T. E., Leblanc, L., & Piñero, J. C. (2010). Recent advances in methyl eugenol and cue-lure technologies for fruit fly detection, monitoring, and control in Hawaii. Vitam Horm, 83, 575–595. 10.1016/s0083-6729(10)83023-7

Wu, H., Appel, A. G., & Hu, X. P. (2013). Instar Determination of Blaptica dubia (Blattodea: Blaberidae) using Gaussian Mixture Models. Annals of the Entomological Society of America, 106(3), 323–328. 10.1603/an12131

Ye, H., & Liu, J. (2007). Population dynamics of oriental fruit fly Bactrocera dorsalis (Diptera: Tephritidae) in Xishuangbanna, Yunnan Province, China. Frontiers of Agriculture in China, 1(1), 76–80. 10.1007/s11703-007-0014-y

